# Dynamic lid domain of *Chloroflexus aurantiacus* Malonyl-CoA Reductase controls the reaction

**DOI:** 10.1101/2023.03.21.533589

**Authors:** Burak V. Kabasakal, Charles A. R. Cotton, James W. Murray

## Abstract

Malonyl-Coenzyme A Reductase (MCR) in *Chloroflexus aurantiacus*, a characteristic enzyme of the 3-hydroxypropionate (3-HP) cycle, catalyses the reduction of malonyl-CoA to 3-HP. MCR is a bi-functional enzyme; in the first step, malonyl-CoA is reduced to the free intermediate malonate semialdehyde by the C-terminal region of MCR, and further reduced to 3-HP by the N-terminal region of MCR. Here we present the crystal structures of both N-terminal and C-terminal regions of the split MCR from *C. aurantiacus*. A catalytic mechanism is suggested by ligand and substrate bound structures, and structural and kinetic studies of MCR variants. Both MCR structures reveal one catalytic, and one non-catalytic SDR (short chain dehydrogenase/reductase) domain. C-terminal MCR has a lid domain which undergoes a conformational change and controls the reaction. In the proposed mechanism of the C-terminal MCR, the conversion of malonyl-CoA to malonate semialdehyde is based on the reduction of malonyl-CoA by NADPH, followed by the decomposition of the hemithioacetal to produce malonate semialdehyde and coenzyme A. Conserved arginines, Arg734 and Arg773 are proposed to play key roles in the mechanism and conserved Ser719, and Tyr737 are other essential residues forming an oxyanion hole for the substrate intermediates.

## Introduction

Improving the efficiency of carbon fixation is one of the goals of crop production improvement to meet growing global food needs^1,2^. Engineering enzymes involved in carbon fixation is of interest not only to increase photosynthetic efficiency but also for biotechnological applications such as production of fuels and organic molecules ^4,5^. The Calvin-Benson cycle has been the classic target for enhancing activity and increasing photosynthetic efficiency, but there are several other carbon fixation pathways^6^. Among the alternative carbon fixation pathways, the 3-hydroxypropionate (3-HP) bi-cycle of *Chloroflexus aurantiacus* stands out, as none of its enzymes are oxygen sensitive and both of the carboxylating enzymes of the cycle, acetyl-CoA carboxylase and propionyl-CoA carboxylase, use bicarbonate rather than carbon dioxide, so there is no oxygenase reaction^7^. 3-HP, an intermediate of the cycle, is a building block chemical and a bioplastic precursor^8,9^. The route from acetyl-CoA to 3-HP is of particular biotechnological interest as 3-HP is an industrial precursor chemical. In the 3-HP bi-cycle route, acetyl-CoA is carboxylated to malonyl-CoA, then 3-HP is generated from malonyl-CoA by malonyl-CoA reductase (MCR) (Fig. 1). MCR can also act as a tartronyl-CoA reductase, producing glycerate and has been used as part of an engineered carbon fixing module^10^.The structure of this, and related enzymes will give better understanding of how they may be engineered, and their specificities may be altered.

**Figure 1.**
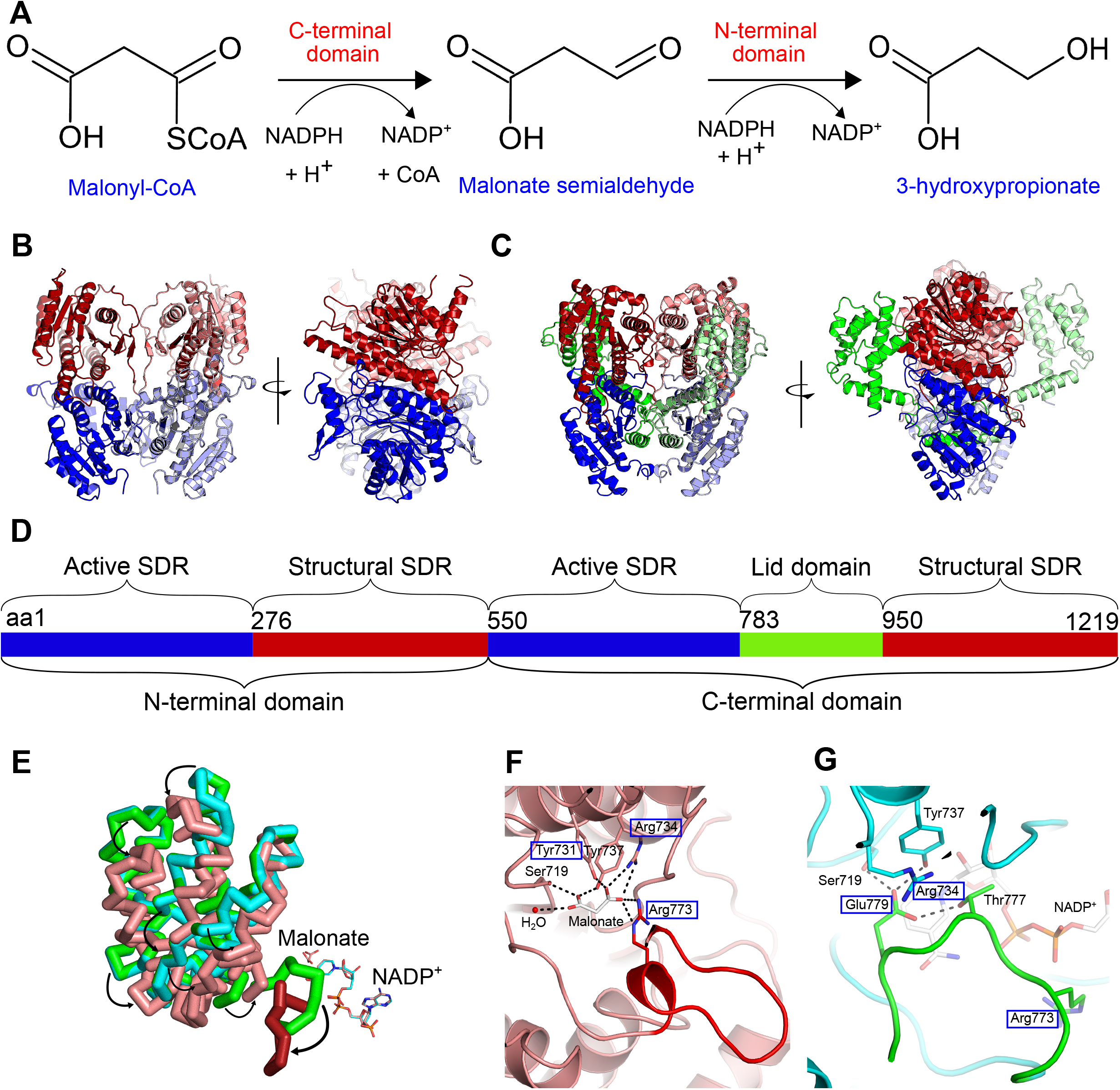
**A. MCR reaction steps. B-C, Orthogonal views of the dimeric N-terminal and C-terminal domains of malonyl-CoA reductase structures, respectively**. Both revealed the predicted SDR domains (blue, light blue), and an additional SDR domain (red, salmon), MCR-C also has a “lid” domain (green, light green). **D. Domain structure of the MCR protein. E, Superposition of lid domains in apo, NADP, and NADP/malonate-bound MCR-C structures in green, turquoise, and salmon.** The loop formations are shown in green and red in NADP-bound, and NADP/malonate-bound structures, respectively. The “lid” domain does not undergo a conformational change in the NADP-bound MCR-C (green), whereas it moves towards the active site (shown with black arrows) in the NADP/malonate bound MCR-C (salmon). NADP has the same conformation in both ligand bound structures. **F, Residues involved in loop formation and catalysis in the NADP/malonate-bound MCR-C.** The loop is shown in red. NADP is not shown for clarity. **G. Residues involved in loop formation in the NADP-bound MCR-C.**The loop is shown in green. The residues we mutated are shown in blue squares.

MCR from *Chloroflexus aurantiacus* (*Ca*MCR) is a 1219 amino acid homodimeric bifunctional enzyme (monomer size ~145 kDa). *Ca*MCR catalyses the conversion of malonyl-CoA to 3-HP in two steps^11^ (Fig. 1A). In the first step, malonyl-CoA is reduced to the free intermediate malonate semialdehyde by the C-terminal domain of MCR, and further reduced to 3-HP by the N-terminal domain of MCR.

The split of *Ca*MCR into two polypeptide chains resulted in increased total enzyme activity when the N-terminal and C-terminal halves were expressed separately^12^. Structures of MCR N-terminal and C-terminal regions from *Porphyrobacter dokdonensis* were recently published^13^. We have solved the homologous regions from *Chloroflexus aurantiacus* MCR, with additional substrate-bound states, and kinetic and structural data from mutants, giving more insight into the mechanism.

## Results

### C-terminal MCR structure has a supernumerary SDR domain and a novel lid region

We solved the crystal structure of the C-terminal *Ca*MCR region (MCR-C). Our C2 space group crystal form contained a monomer, which expands to the dimer by crystallographic symmetry. MCR-C has two SDR (short chain dehydrogenase) domains^14^, with a novel alpha-helical “lid” domain between them. The first SDR domain (Fig. 1C blue) has the TGXXX[AG]XG cofactor binding YXXXK active site motifs^14,15^ between residues 578-585 and 737-741 respectively, corresponding to NADPH and malonyl-CoA binding sites. Neither of these motifs were found in the second domain (Fig. 1C red), which is unlikely to be catalytic. The lid domain has no significant structure matches in the PDB except to the equivalent region in the MCR-C from *P. dokdonensis*. Overall, our apo MCR-C is similar to the *P. dokdonensis* structure^16^, with 52% sequence identity and 2.0 Å RMSD over the length of the apo-proteins. The full length *Ca*MCR has 45% sequence identity with *P. dokdonensis* MCR.

### Ligand binding causes conformational changes in the MCR-C structure

We obtained two ligand-bound MCR-C structures; NADP-bound, and NADP and malonate-bound MCR-C. We co-crystallised MCR-C with NADP^+^ to get the NADP^+^ bound structure. Co-crystallisation of MCR-C with NADPH gave an NADP-bound MCR-C structure with an additional adventitiously bound small molecule which we identified as malonate. We propose that this malonate is the oxidation product of malonate semialdehyde, trapped in the closed product bound conformation of the enzyme.

NADP^+^ has an S-shaped conformation in the MCR-C structure (Fig. 1E). In the apo crystal structure of MCR-C, there was no interpretable electron density for residues 771 to 783, which is between the first SDR domain and lid domains. But in the NADP-bound, and NADP/malonate-bound MCR-C structures, residues 771 to 783 formed a loop adjacent to the NADP binding site, in a different conformation in each structure (Fig. 1E, F, G). In the NADP/malonate-bound structure, the loop formed a 13 Å-long hairpin with the two ends pointing towards the active site. The ordering of the hairpin loop brings Arg773 to the active site, where the side chain forms a hydrogen bond to malonate in our structure (Fig. 1F).

In the ligand bound structures, the lid domain is in a “closed” conformation, closer to the SDR domains. In the apo structure it is “open”, rotating 15 degrees around an axis in the middle of the domain, and moving Gly782 in the hairpin loop by 8.7 Å towards the active site. The lid domain interacts with the second SDR domain, which can be assigned a possible function as an anchor for the domain motion.

In the NADP-bound MCR-C structure, NADP has the same S-shaped conformation as in the NADP/malonate-bound structure. The active site loop is mainly ordered, residues 782-784 are missing, but is in a different conformation, where Arg773 is pointed away from the active site. The lid domain is in the open conformation, similar to the apo structure. Therefore we think that this structure is not in a catalytically competent conformation (Fig 1G).

Although the MCR-C structure is very similar to its homolog from *P. dokdonensis* including the bound NADP (Fig. S3), the observation of the conformational change in the “lid” domain is specific to *C. aurantiacus* MCR-C. In the Pd MCR-C structures, either apo or bound to NADP^+^ or coenzyme A (CoA), the active site loop is ordered, and the lid domain is in what we call the closed state. It is possible that the CaMCR-C crystal form allows us to observe these conformational changes.

In the NADP binding site, the side chains of Tyr737 and Lys741 are hydrogen-bonded to the ribose ring carrying nicotinamide along with the main chain carboxyl groups of Asn666 and Val717. Asp639 makes a hydrogen bond with the adenine moiety, and Ser581 interacts with the ribose ring carrying adenine. The phosphate groups interact with the main chain amide and carbonyl groups, and water molecules (Fig. S4). Compared to the apo structure, Arg604 and Lys608 move towards the NADP. Arg604 makes hydrogen bonds with NADP in both NADP-bound and NADP/malonate-bound structures whereas Lys608 is hydrogen-bonded with NADP only in the malonate-bound structure (Fig. S4). Thr777 and Glu779 appear near the NADPH binding site in the NADP-bound structure (Fig. 1G) whereas Arg773 is ordered and interacts with not only the pyrophosphate of NADP (not shown in the figure) but also malonate in the NADP/malonate-bound structure (Fig. 1F). Arg773 is stabilised by a hydrogen bond to the Gly668 carbonyl. Although the residues in the loop, Thr777 and Glu779, do not interact with NADP in the NADP-bound structure, other interactions seem to help the loop keep in place. The side chain of Thr777 is in close contact with the Glu779 side chain, and Glu779 is hydrogen bonded with both Tyr737 and Ser719 (Fig. 1G). Thr777 and Glu779 are on the turn of the hairpin loop away from the active site in the malonate bound structure, so they are unlikely to be involved in catalysis.

### Malonyl-CoA and NADPH binding cooperate

Several attempts were made to solve the structure of MCR-C bound to different substrates by cocrystallization, however no CoA substrate including malonyl-CoA could be located.

To investigate whether added malonate ions have any effect on conformation as malonyl-CoA, MCR-C was co-crystallised or soaked with various concentrations of sodium malonate (0.5 – 20 mM) with or without NADPH (0.5 mM – 2 mM). However, no malonate was found near the active site. The effect of malonate ions on the MCR-C reaction was also investigated. Malonate did not stimulate or inhibit the reaction. These results support two hypotheses: (i) the malonate ion in the active site comes from malonyl-CoA reduction to malonic semialdehyde, followed by oxidation to malonate, (ii) free malonate is not an intermediate of the MCR-C catalysis. Thermodynamic calculations suggest that NADPH cannot reduce malonate to malonic semialdehyde, as the Gibbs free energy of the malonate to malonate semialdehyde conversion is 36.1 kJ/mol at physiological conditions ^17^.

The malonate ion is tightly bound in the active site making hydrogen bonds with polar residues such as Ser719, Tyr731, Arg734, Tyr737, and Arg773. The previously mentioned hydrogen bond of Arg773 to malonate suggests that the binding of malonyl-CoA forms this hydrogen bond, ordering Arg773 and the active site loop, pulling the lid domain into the closed conformation required for catalysis (Fig. 1E, 1F).

### Site-directed mutagenesis reveals catalytic residues

The mutations in this study were designed to probe the putative malonyl-CoA binding site, and to determine the residues involved in the catalysis. We could measure no activity towards malonyl-CoA for any of our six MCR-C variants (Arg734Gln, Arg734Ala, Arg773Gln, Arg773Ala, Tyr731Ala, and Glu779Trp) (Fig. S5). Neither NADP nor malonyl-CoA could be found in the crystal structures of variants. These results suggest that Arg734, Arg773, Tyr731, and Glu779 are essential for NADPH binding and catalytic activity.

We propose that Arg734 is an essential residue for catalytic activity. It is hydrogen bonded to malonate in the NADP/malonate-bound structure. It may function as a base to produce malonate semialdehyde from the hemithioacetal formed by NADPH reduction. Given this hypothesis, Arg734 was mutated to Ala and Gln. Gln734 in Arg734Gln was predicted to still interact with malonyl-CoA/malonate. The Arg734Gln variant did not show any activity. We therefore find the acid-base properties of Arg734 are essential for MCR-C activity. Unfortunately, there was no structural information on any of the Arg734 variants as no crystals could be obtained. The mutation of Arg to Ala was predicted to eliminate the interaction between malonyl-CoA and Arg734. The Arg734Ala variant was also completely inactive, indicating the interaction between Arg734 and malonyl-CoA was vital.

Arg773 was hydrogen bonded to both malonate and NADP in the NADP and malonate-bound structure. These strong interactions between Arg773 and the active site may keep the loop stable and in place. The Arg773Ala and Arg773Gln mutations completely eliminated MCR-C activity. Both Arg773Ala and Arg773Gln variant structures were determined. Neither malonyl-CoA/malonate nor NADP was obtained bound in the crystal structures. In the Arg773Ala structure, the mutant residue was not visible and the lid domain was in the open conformation. In the Arg773Gln structure, the loop was partially ordered and the mutant Gln773 residue was visible, but not in a position to hydrogen bond to the malonate (Fig. S7). The Gln773 C_α_ atom was 4.4 Å from the Arg773 C_α_. In Arg773Gln, the lid domain was in a partially closed conformation, intermediate between the apo and malonate bound structures. The Arg773Ala and Arg773Gln mutations showed that Arg773 is essential for MCR-C activity. It is the key residue for not only conformational change and loop formation, which is required for malonyl-CoA and NADPH binding, but also for malonyl-CoA conversion.

Tyr731 was mutated to Ala to investigate its importance in activity. In the NADP/malonate-bound MCR-C structure, Tyr731 was hydrogen bonded to the malonate ion. The Tyr731Ala mutant structure had no substrates bound and a disordered loop (Fig. S7). As in Arg773Gln, the lid domain was in the partially closed conformation (Table S4). Malonyl-CoA and NADPH may have bound only weakly, as no activity of the Tyr731Ala mutant could be measured.

Finally, the effect of the mutation Glu779Trp was investigated. Glu779 was ordered in the NADP-bound structure and found in same position as malonate in the NADP/malonate-bound structure. On malonyl-CoA binding, the Glu779 would move to allow the malonyl moiety to bind. According to this hypothesis, relocation of Glu779 and malonyl-CoA binding would be eliminated by replacing Glu779 with a large residue such as Trp. The Glu779Trp variant did not show any measurable activity. The Glu779Trp structure has an “open” state and was similar to the apo MCR-C structure (RMSD = 0.25 Å) as expected. Trp779 was not seen in the density (Fig. S8).

### Proposed reaction mechanism for MCR-C

We propose the initial state of the enzyme is in the “open” conformation with a flexible loop, as in the NADP-bound MCR-C structure. NADPH binding occurs before malonyl-CoA binding. Once malonyl-CoA enters the enzyme, binding of Arg773 and other residues to the substrate orders the active site loop and pulls the lid domain into the closed conformation.

The reduction of malonyl-CoA by NAPDH and proton donation from Arg734 form a hemithioacetal. After the Arg734-catalyzed decomposition of the hemithioacetal to produce malonate semialdehyde and the CoA thiolate anion, free CoA leaves the enzyme after the protonation of Arg773. Once the malonate semialdehyde site is unoccupied, the loop returns its initial conformation, and the lid domain moves away from the active site to the open state. Finally, NADP^+^ is released from the enzyme (Fig. 2).

**Figure 2.**
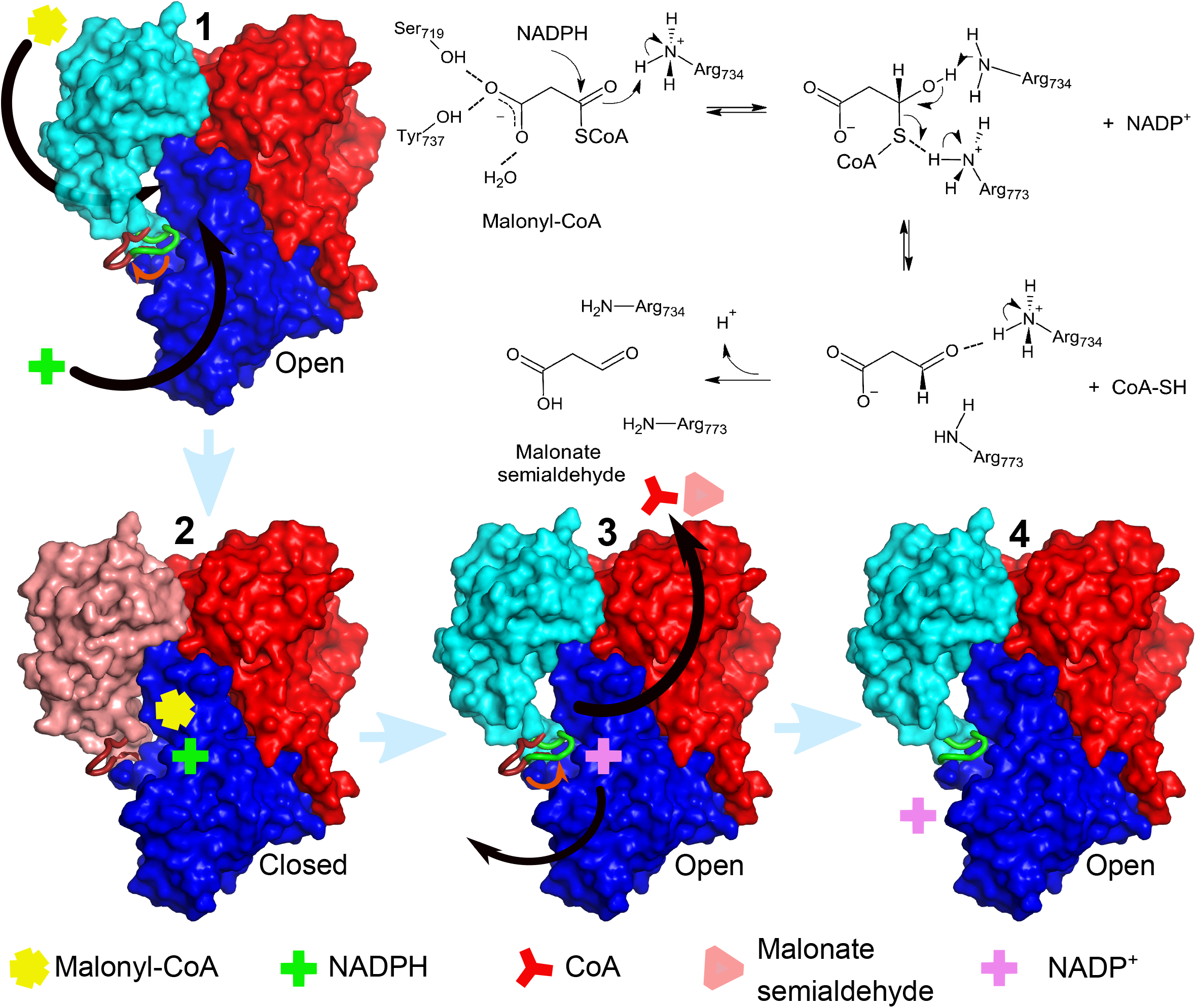
Proposed mechanism for the MCR-C. The diagram shows the proposed states of MCR-C during the reaction. **1.** MCR-C is in the “open” state with a loop (green) positioned towards the active site. **2.** Once malonyl CoA binds, the “lid” domain moves towards the active site, and the loop forms a hairpin (red). NADPH binding occurs with or before malonyl-CoA binding. **3.** NADPH reduces malonyl-CoA, malonyl-CoA is then converted to malonate semialdehyde, and free CoA is released from the enzyme. **4.** Once malonate semialdehyde leaves the malonate binding site, the “lid” domain moves back to the initial state, and the loop positions towards the active site again. NADP^+^ leaves the enzyme, and MCR-C returns the initial state. Both loop conformations (green for the open state, red for the closed state) are shown to highlight conformational changes (shown with orange arrows). The “lid” domain is shown in turquoise and salmon for open and closed states, respectively. **Inset. Proposed reaction steps.** NAPDH reduction of malonyl CoA and proton donation from Arg734 produce the hemithioacetal. Aldehyde bond formation occurs with Arg734-assisted catalysis and Arg773 protonation of the thiol bond of hemithioacetal to release CoA. Ser719 and Tyr737 stabilize malonyl moiety by forming an oxyanion hole.

Based on the structures, and enzyme kinetics, we propose the mechanism for MCR-C to convert malonyl-CoA to malonate semialdehyde is based on the reduction of malonyl-CoA by NADPH to a hemithioacetal, followed by the base-catalyzed aldehyde formation, and release of free CoA simultaneously (Fig. 2 Inset). The proposed mechanism is similar to that of 3-Hydroxy-3-methylglutaryl coenzyme A reductase (HMGR) from *Pseudomonas mevalonii*^18^. HMGR has a disordered flap domain which becomes ordered over the active site when NAD^+^and HMG-CoA bind. A lysine residue makes a hydrogen bond with the thioester carbonyl of the substrate in HMGR crystal structure^18^. In our proposed mechanism, NAPDH reduces malonyl CoA and Arg734 donates protons to the carbonyl of malonyl-CoA to form a hemithioacetal. Malonate semialdehyde from the hemithioacetal is produced after the aldehyde bond formation, and the free CoA is formed by the protonation from Arg773 subsequently. Malonate formation is proposed to be favorable after the malonate semialdehyde formation, as the intermediate product, malonate semialdehyde is not stable and may be converted to malonate by hydrolysis. The malonate (and possibly malonate semialdehyde during the reaction) is stabilized by an oxyanion hole formed by Ser719, Tyr737, and a water molecule found in the crystal structure. Malonate semialdehyde is further reduced to 3-hydroxypropionate by the N-terminal domain of MCR (MCR-N).

### MCR-N structure also reveals an additional SDR domain

The 2.8 Å X-ray crystal structure of MCR-N (Fig. 1A) contains a biological dimer. Each monomer has two SDR domains (one predicted and one additional) were found, like the MCR-C, as in the *P. dokdonensis* N-terminal malonyl-CoA reductase (Fig. S8)^13^. The second SDR domain had a mean B factor of 97, compared with 82 for the first domain, less clear electron density and two unmodelled loops at residues 442 and 457. The dimer partners are connected by a disulfide bond between Cys238 on each monomer, although this residue is not conserved.

The first SDR domain (1-275aa) has a nucleotide binding GXGXXG motif. The putative NADPH binding site is the TG_16_GAGNIG_22_ region. There is a Y_171_VTPK_175_ motif near the di-nucleotide binding domain, which is proposed to be the active site. There is a cleft between TG_16_GAGNIG_22_ and Y_171_VTPK_175_ regions, where NADP may bind in the *C. aurantiacus* structure (Fig. 3). In the MCR-C, which has similar motifs for dinucleotide and substrate binding, there is a cleft where NADP and malonate are bound. The structural alignment of MCR-C and MCR-N SDR domains revealed that these domains including cleft regions are structurally similar (RMSD of 1.5 Å over 141 residues). In MCR-C malonate is hydrogen bonded to the side chains of residues Ser719, Tyr731, Arg734, Tyr737 and Arg773. Arg773 could be equivalent to Arg208 in MCR-N, although both in our structures and the PdMCR it is not pointing into the active site cavity. The other malonate binding residues are conserved in MCR-N as Thr158, Tyr165, Arg169 and Tyr171. We speculate that these residues form the malonate semialdehyde binding site. The only acidic residue near the putative substrate binding site is Glu164, however it is positioned away from the cleft where malonate semialdehyde may bind (Fig. 3B).

**Figure 3.**
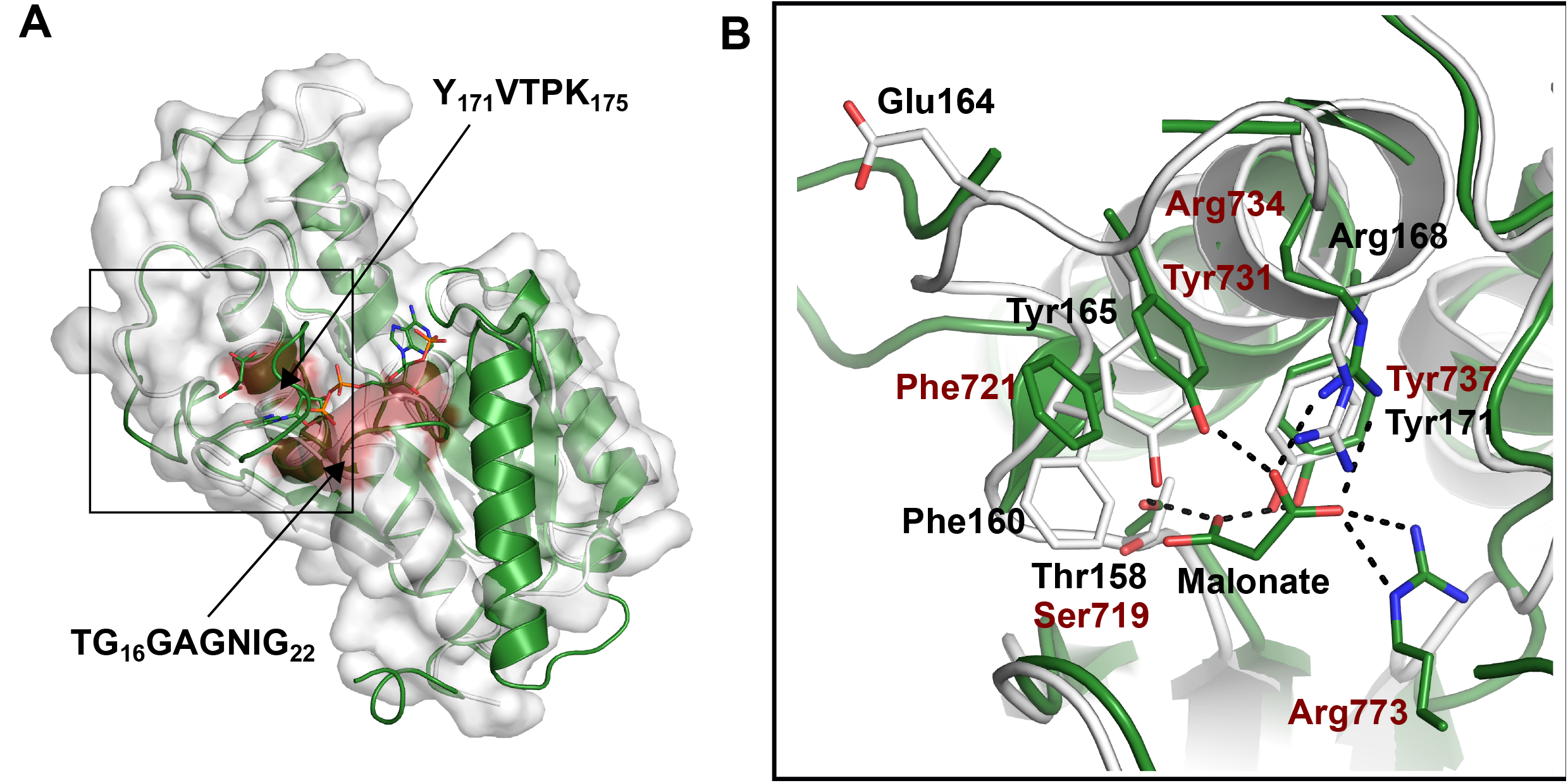
Putative active site of the MCR-N. A. Surface representation of the putative catalytic SDR domain of the MCR-N (white). Y_171_VTPK_175_ and TG_16_GAGNIG_22_ motifs are shown in red. The cleft is located between these motifs, where NADPH and substrate may bind. The green cartoon model represents the catalytic Rossman fold domain of the MCR-C superposed. Malonate and NADP^+^ are shown in green sticks. **B. Close view of the putative substrate binding site of the MCR-N.** Putative active residues, and Glu164 are shown in white sticks (black labels). Superposing MCR-C residues (red labels) and bound malonate are shown in green sticks.

### Prediction of the full-length MCR structure

After we solved the crystal structures, Alphafold2 became available and so we predicted the structure of the full-length MCR dimer^19^ (Fig 4). The MCR-C lid domain is in the closed conformation in the predicted structure. The crystal structures of MCR-C and MCR-N are a good fit to the predicted structure (RMSDs = 0.84 and 0.63 Å, respectively). The predicted structure revealed an elongated dimer as in the *P. dokdonensis* MCR SAXS model^13^. In the *Pd*MCR SAXS model the faces are parallel, but in the prediction, they are twisted by around 90 degrees, however the relative orientation of the MCR-C and MCR-N is less confident, according to the PAE (predicted alignment error matrix, Figure S15).

**Figure 4.**
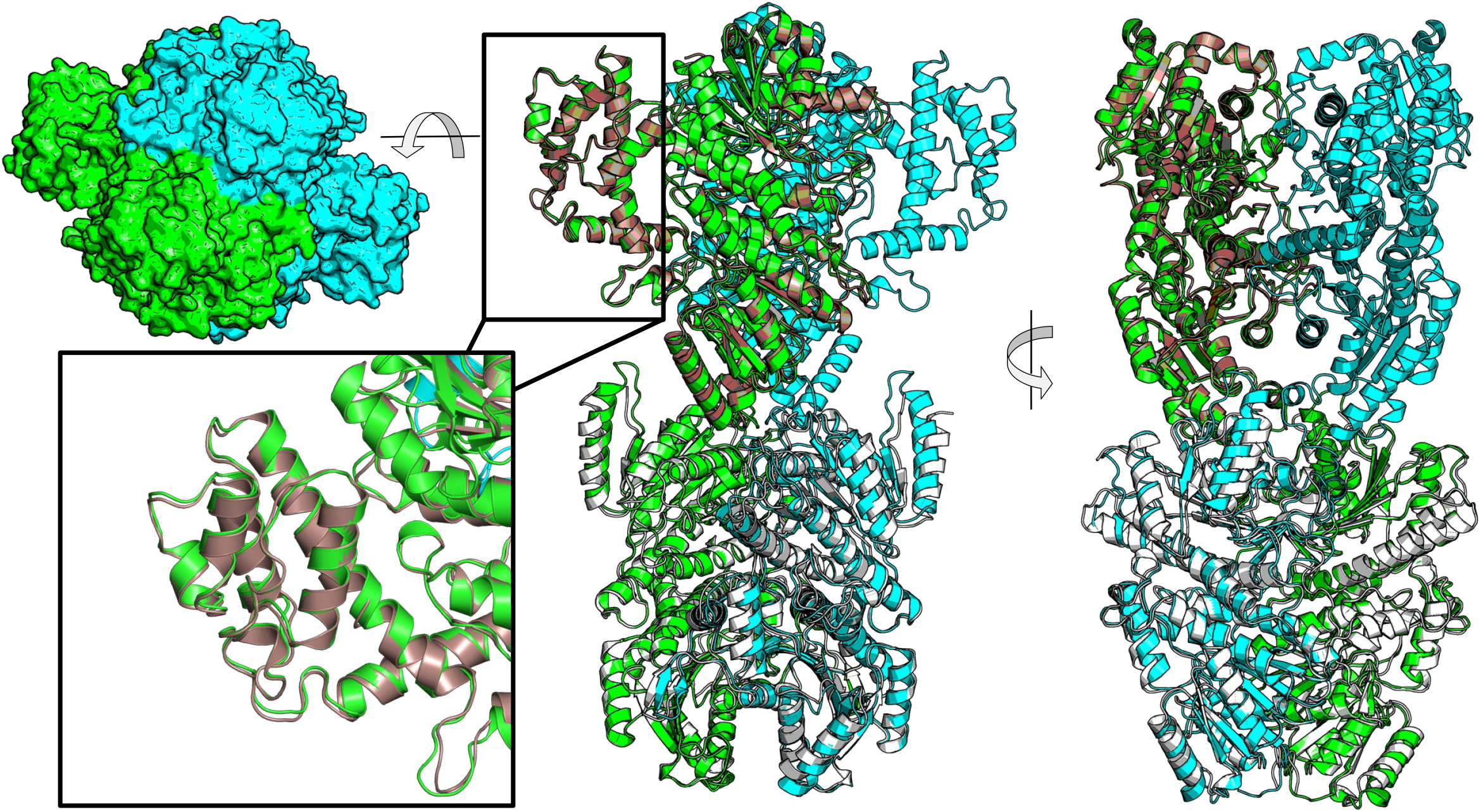
Orthogonal views of the predicted structure of the full-length MCR and comparison with MCR-N and MCR-C crystal structures. The dimeric full-length MCR is shown in turquoise, and green (the top view is shown in surface representation for clarity). MCR-N and MCR-C are shown in white, and brown, respectively. NADP/malonate bound MCR-C crystal structure is used for superposition to highlight the similarity (black box) (NADP and malonate are not shown).

### MCR Activity assays and enzyme kinetics

The activities of MCR-C and MCR-N were measured using 0.1 mM malonyl-CoA, and 10 μg of MCR-C at room temperature (~25 °C) as 1.85 ± 0.22 μmol min^-1^ mg^-1^ MCR-C protein and 0.37 ± 0.04 μmol min^-1^ mg^-1^ MCR-N protein (n=5), respectively (Tables S9, S10). The K_m_ of MCR-C was 0.0174 ± 0.0030 mM (2≤n≤5) for malonyl-CoA, and the K_m_ of MCR-N was determined as 1.27 ± 0.23 mM (2≤n≤5) for malonate semialdehyde. The MCR-C was ~5 times faster than the MCR-N at MCR-C substrate saturating conditions (0.1 mM malonyl-CoA, K_m__MCR-C=0.017 mM, K_m__MCR-N=1.27 mM). However, the turnover rate of MCR-N is nearly 2-fold higher than MCR-C but the k_cat_/K_m_ is very low compared to that of MCR-C due to the low substrate affinity (K_m_ 1.27 mM). MCR-N has consequently a lower catalytic efficiency than MCR-C (Table 1). The activity of MCR-C towards malonate ions was investigated with 0.2 mM and 0.4 mM sodium malonate. MCR-C showed no detectable activity in the presence of malonate ions even in excess (1.5-fold) compared to NADPH. Malonate was also investigated as a possible competitive inhibitor. Malonate was added to the reaction medium containing 0.1 M malonyl-CoA. The inhibition was investigated for malonate at 0.1 mM, 0.5 mM, and 1 mM, and no inhibition was found. The activity of MCR-C variants were investigated using the same reaction conditions as WT. MCR-C reactions were measured for R734A, R734Q, R773A, R773Q, Y731A, and E779W variants. None of them showed any measurable activity towards malonyl-CoA (Fig. S5).

**Table 1.** Kinetic parameters of MCR-C and MCR-N

| | $K_m$ (mM) | $k_{cat}$ (1/s) | $k_{cat}/K_m$ (1/mM.s) |
| --- | --- | --- | --- |
| MCR-C | 0.017 | 2.61 | 150.1 |
| MCR-N | 1.270 | 4.80 | 3.8 |

## Discussion

MCR-C has no activity towards malonate, and we could not solve a structure with malonate added to the crystallization mixture. The affinity of the open structure for malonate appears to be low. Therefore, our malonate bound structure probably represents a malonate semialdehyde product complex that has been oxidised to malonate in the crystal. In the archaeal MCR, both NADPH and malonyl-CoA must bind sequentially as they share a binding site ^20^. In MCR-C it appears that NADPH and malonyl-CoA are both bound before the reaction, and the productive complex is formed by the closure of the lid domain.

The MCR-C and MCR-N dimers each have four SDR domains, forming an approximate D2 tetramer, corresponding to the homotetrameric structure of single domain SDRs (Fig S15). The MCR-C and MCR-N regions probably arose by duplication of an ancestral SDR domain. One domain then lost its catalytic function and became a supernumerary domain. Other enzymes are known to have supernumerary domains, for example the non-catalytic domains of polyketide synthases^21^. There are many known examples of pseudoenzymes, which maintain an enzyme-like structure, but have lost catalytic activity. The non-catalytic domains of MCR-C and MCR-N are conserved in many species, implying that they are required for activity, even if not catalytic. With the publication of comprehensive databases of protein structure predictions^22^, more non-catalytic enzyme domains will come to light.

In *E. coli* cells with both MCR-C and MCR-N present, the MCR-C activity appeared to be limiting^12^. However, in our work, the MCR-C was measured to be five times faster than MCR-N when MCR-C was saturated. The explanation is likely to be that in the earlier work^23^ the MCR-N was present in excess, making MCR-C limiting. Here we measured the K_m_ and k_cat_ of both enzyme activities separately. Given the K_m_ and k_cat_ values we may characterise MCR-C as “slower but tightly binding” to the substrate, and MCR-N as “fast, but loosely binding”. In the *Chloroflexus* cell, the MCR-C and MCR-N are necessarily present in equimolar amounts. There is likely some substrate channelling, increasing the effective concentration of the malonate semialdehyde intermediate which mitigates the higher K_m_ of MCR-N.

Liu *et al*. (2013) measured the activities of double and single site variants of MCR-C (Y737G/K741G and G579R), and showed that Y737, K741, and G579 are essential for activity^12^. In our structure, Lys741 is hydrogen bonded to NADP, and Tyr737 is hydrogen bonded with both NADP and malonate. Their contribution may be related to NADPH/malonyl-CoA binding, and catalytic activity. Gly579 is part of the NADPH binding pocket. We designed the mutations to determine the active site residues. The structures of the Arg773Ala, Arg773Gln, and Tyr731Ala variants showed that malonyl-CoA binding causes the movement of the lid domain towards the active site. Arg773 is the main residue involved in the movement due to its interaction with both malonate and NADP. These interactions were disrupted in the R773A variant, and it revealed a structure in the “open” state as in the apo MCR-C. However, both R773Q and Y731A structures were in the “partially closed” state as in the malonate - bound MCR-C. The extent of lid movement was higher in the Y731A variant than in the R773Q variant, suggesting that Arg773 functions in the conformational change. In the R773Q variant, the non-complete movement of the lid domain may be because of weak interactions with either malonyl-CoA or NADPH. Both Arg773 and Tyr731 are required for malonyl-CoA binding and to stabilise the intermediate malonate.

None of our MCR-C mutants had any detectable activity. Mutants in the *P. dokdonensis* MCR^16^ made by Son *et al*. (2019) were Y731A equivalent (Y738) with ~10% wild-type activity, and R773A equivalent (R789A), with ~5% wild-type activity. These results show that these are important residues for activity.

Glu779 is located in the malonate site in the NADP-bound structure. As the disordered hairpin region was assumed to have the same conformation in the apo enzyme as in the NADP-bound structure, Glu779 would be expected to move towards solvent to allow the malonyl-CoA to bind the malonate site. The conformational change in the hairpin would occur at the same time as the lid movement. Based on this hypothesis, the entrance and binding of malonyl-CoA to the malonate binding site was aimed to be prevented by substituting Glu779 with a large residue Trp. The Glu779Trp variant was not active, and its structure was in the “open” state, suggesting that malonyl-CoA binding did not occur.

The additional SDR domain (276-549aa) of the MCR-N is possibly a non-catalytic, regulatory domain. As in the MCR-C, this additional SDR domain (the other half of the MCR-N) has neither cofactor binding motif (TGXXX[AG]XG) nor active site motif (YXXXK). Consequently, we hypothesise that NADPH binds only the putative NADPH binding site in the first predicted SDR domain (1-275aa) as no ligand bound MCR-N structure could be obtained. The second step of the MCR reaction, reduction of malonate semialdehyde to 3-HP, probably occurs in this domain.

The sequence alignment of MCR-N with other SDR proteins showed that both cofactor binding motif (TGXXX[AG]XG) and active site motif (YXXXK) are conserved in Chloroflexi, Roseiflexi, Proteobacteria, Alphaproteobacteria, and Gammaproteobacteria (Fig. S10). There are conserved residues, Tyr165, Arg168, and Tyr171 near the active site motif, as in the MCR-C. They may be catalytically important residues as they are predicted to bind malonate semialdehyde, by analogy with the MCR-C active site (Fig. 3).

The conversion of malonate semialdehyde to 3-hydroxypropionate requires further reduction of the carbonyl bond of the aldehyde with a base-catalyzed mechanism. In MCR-N, Glu164 is the only acidic residue that could play a nucleophilic role. It is conserved in Chloroflexi species and is substituted with an Asp in Roseiflexi (Fig. S10). However, it is positioned towards the outside, and away from the cleft that malonate semialdehyde may bind (Fig. 3B). Therefore Glu164 is probably not a catalytic residue, and MCR-N has a similar mechanism to MCR-C, in which no acidic residue is involved, and Arg168 is the main catalytic residue.

Structural, mutagenesis, and kinetic studies of MCR-C and MCR-N gave insights into MCR, and the mechanism of the reaction generating 3-HP from malonyl-CoA. First of all, MCR from *C. aurantiacus* is twice as large as necessary for activity. The dynamic lid domain changes its conformation during the reaction, and controls the mechanism. Many conserved residues are involved in catalytic activity, stabilization of the intermediate and conformational change. Future work is required to understand whether the conformational change is the case for the full native MCR, also for other homologs. The results of this study are valuable for not only biological 3-HP production but also future biotechnological studies, and contribute to strategies for CO_2_ use to minimise the effects of global climate change.

## Materials and Methods

### Cloning, purification, and crystallisation

*Chloroflexus aurantiacus* J-10-fl Malonyl-CoA reductase (*Caur_2614*) N-terminal region (residues 1-549) construct and Malonyl-CoA reductase C-terminal region (residues 550-1219) construct were cloned into a modified pRSET-A vector ^24^ by Gibson assembly ^25^. Primers (Invitrogen) used for the pRSET-A vector were 5’-GGATCCACGCGGAACCAGACCATGATGATGATGATGATGAGAACCCCGCAT-3’ (forward), 5’-AAGCTTGATCCGGCTGCTAACAAAGCCCGAAAGGAAGCTGAGTTGGC-3’ (reverse), for MCR-N were 5’-CATCATCATCATCATCATGGTCTGGTTCCGCGTGGATCCATGAGCGGAACAGGACGA CTG-3’ (forward), 5’CTTCGGGCTTTGTTAGCAGCCGGATCAAGCTTTTAAATGTTGGCAGGGATGTTGAGGG-3’ (reverse), and for MCR-C are 5’-CATCATCATCATCATCATGGTCTGGTTCCGCGTGGATCCAGCGCCACCACCGGCGC ACGC-3’(forward), 5’-GCTTCCTTTCGGGCTTTGTTAGCAGCCGGATCAAGCTTTTACACGGTAATCGCCCG-3’ (reverse). The plasmids were transformed into *E. coli* KRX^26^ for expression. Cells were grown at 18°C for 18 h after induction, harvested, resuspended in 50 mM Tris-HCl, pH 7.9, 150 mM NaCl, and disrupted by sonication. The supernatant was loaded onto a nickel resin affinity column (Generon) and eluted with 300 mM imidazole in 50 mM Tris-HCl pH 7.9, 150 mM NaCl. The His-tag of the eluted protein was cleaved by thrombin (Sigma), incubated at 4 °C overnight. The cleaved protein was then purified by gel filtration on a Superdex 200 HiLoad 16/600 column with buffer 50 mM Tris-HCl, pH 7.9, 150 mM NaCl. The proteins were concentrated to 10-15 mg/ml for crystallisation. Selenomethionine based expression of MCR-N and MCR-C was performed using the Promega selenomethionine protein labelling protocol for KRX cells^27^.

Proteins were crystallised by sitting-drop vapour diffusion against sparse-matrix screens, set up with a Mosquito robot (SPT Labtech), then with manual optimization plates. We found many crystallization conditions for MCR-C, but only three for MCR-N (Table S5, S6). Crystals were cryoprotected in the mother liquor with 30% volume PEG (polyethylene glycol) 400 added.

### Structure determination, refinement, and analysis

X-ray diffraction data of cryoprotected crystals were collected at 100 K at Diamond Light Source, UK. The collected data were processed and scaled with *xia2* using the 3dii setting^29^ or DIALS^30^. The experimental phases for MCR-C were obtained and refined with SHELXD and SHELXE^31^. Model building and manual rebuilding was performed in BUCCANEER^32^ and COOT^33^, and refined with REFMAC5^34^ and phenix.refine^35^. NADP/malonate bound MCR-C and variant MCR-C structures were solved with molecular replacement with PHASER^36^ using the native structure as a model.

The initial model for MCR-N was obtained with CRANK^37,38^ pipeline using the Se-Met and a higher resolution native data set. The model was manually completed in COOT ^33^ and refined using REFMAC5^34^ and phenix.refine^35^ against a higher resolution native dataset. Structures were validated with MolProbity^39^. Data collection and refinement statistics are given in Tables S1-3, and S7. Molecular figures were made with PyMol^40^. AlphaFold^19^ was run through ColabFold^41^

### Site-directed mutagenesis

MCR-C mutants, R734A, R734Q, Y731A, R773A, R773Q, and E779W were generated with the Q5 Site-Directed Mutagenesis Kit (E0554, New England Biolabs). The primers were designed with the NEBaseChanger^®^ tool (Table S8).

### MCR activity and kinetic measurements

MCR-C and MCR-N activities were measured based on NADPH oxidation. The assay mixture (500 μl) contained 100 mM Tris-HCl (pH 7.8), 2 mM MgCl_2_, 0.1 mM malonyl-CoA, 0.4 mM NADPH, and 10 μg of purified MCR-C or MCR-N proteins. The reactions were carried out in 100 μl quartz cuvettes, started by the addition of individual enzymes and incubated at room temperature (~25°C) for 3-10 minutes. MCR-N activity was measured by using MCR-C to generate the malonic semialdehyde substrate *in situ*, then adding MCR-N. The NADPH concentration was monitored at 340 nm (absorption coefficient at 340 nm is 6,300 M^-1^ cm^-1^ provided by the manufacturer), and the enzyme activity was calculated according to the rate of NADPH oxidation (Fig. S13). The blank was the assay buffer, 100 mM Tris-HCl (pH 7.8), 2 mM MgCl_2_. The MCR-C activity was also measured in the presence of malonate (0.2 mM) instead of malonyl-CoA. To investigate the inhibition of malonate on MCR-C activity, the enzyme activity of MCR-C was measured for 0.1 mM malonyl-CoA, and three malonate concentrations (0.1 mM, 0.5 mM, and 1 mM). Activity measurements of variant MCR proteins were performed in the same way with 0.1 mM malonyl-CoA. Reaction rates were determined based on initial rates.

Michaelis-Menten parameters of MCR-C and MCR-N were determined by the measurement of initial reaction rates for various malonyl-CoA concentrations (0.05 mM, 0.075 mM, 0.1 mM. 0.125 mM, and 0.15 mM) with a Lineweaver-Burk plot (Tables S11, S12, Figures S1, S2). For MCR-N, initial rates were measured by adding N-terminal MCR to the reaction media after the completion of MCR-C reactions.

### HPLC analysis of MCR reaction product

The product of the net MCR reaction, 3-hydroxypropionate, was analysed by HPLC (High Performance Liquid Chromatography) using a photodiode array detector (Jasco HPLC Systems). The reactions carried out with both MCR-C and MCR-N, which were expected to produce 3-hydroxypropionate, were terminated by adding 100 μl pure formic acid, and centrifuged at 14,000 g for 20 min at room temperature (VWR Microstar microcentrifuge, UK) to precipitate the denatured proteins. The supernatants were passed through 0.2 micron filters before analysis.

The analyses (100 μl injection volumes) were performed on a Luna^®^ reversed phase C18 100A analytical column (5 μm, 125 x 4.0 mm, Phenomenex, Torrance, CA) at room temperature with a flow rate of 0.6 ml min^-1^. Two solutions were used as the mobile phase; Solution A, 40 mM KH_2_PO_4_, 0.2 % formic acid, pH 4.2 and Solution B, 100 % methanol and Solution A (70:30, v/v). The standards and reaction product were eluted with a 10-min protocol. For the first 3 min, 100 % of Solution B was used, and between 3-10 min, a linear gradient from 0 % to 50 % of Solution A. The standards, malonyl CoA (0.025 mM and 0.050 mM) (malonyl CoA lithium salt, ≥90 % (HPLC grade), Sigma-Aldrich) and 3-hydroxypropionic acid (18 mM and 36 mM) (Tokyo Chemical Industry Limited, UK) were monitored at 260 and 210 nm, respectively. Their retention times were 4.0 min, and 4.1 min, and the separation was based on the absorption at different wavelengths due to close retention times. The reaction product was monitored at both wavelengths, it had absorption only at 210 nm. A mixture (50:50 v/v) of Solution A and Solution B was used as a blank and monitored at both 260 and 210 nm (Fig. S14).

## Data availability

X-ray crystal structure coordinates and structure factors have been deposited in the PDB, accession codes for the apo MCR-C, MCR-N, NADP-bound MCR-C, NADP and malonate bound MCR-C, Y731A MCR-C, R773A MCR-C, R773Q MCR-C, E779W MCR-C are 8A30, 8AEW, 8A7S, 8A8T, 8AER, 8AEO, 8AEQ, and 8AET respectively.

## Supporting information

This article contains supporting information

## Conflict of interest

The authors declare no competing interests.

## Acknowledgements

We thank Diamond Light Source for X-ray beam time (proposal mx12579) and the staff of beamlines I03, I04 and I04-1 for assistance with crystal testing and data collection. The crystallization facility at Imperial College was funded by the BBSRC (BB/D524840/1) and the Wellcome Trust (202926/Z/16/Z). We acknowledge Imperial College Research Computing Service, DOI: 10.14469/hpc/2232 for AlphaFold calculations.

## Funding information

BVK is supported by a TUBİTAK BIDEB 2232 Fellowship (Project No. 118C225). CARC was supported by a BBSRC Doctoral Training Programme grant (BB/F017324/1).

## Notes

### Competing Interest Statement

The authors have declared no competing interest.

