## Supplementary Files for "Dynamic lid domain of *Chloroflexus aurantiacus* Malonyl-CoA Reductase controls the reaction"

**Table S1. Data collection and refinement statistics for C-terminal Malonyl-CoA reductase (MCR-C) structure.** Values in brackets refer to high resolution shell.

| Data collection | Se-Met MCR-C | Native MCR-C | Refinement | Final MCR-C PDB: 8A30 |
|---|---|---|---|---|
| **Crystallisation condition** | 90 mM Sodium cacodylate, pH 6.5, 1.26 M Sodium acetate trihydrate | 0.8 M Succinic acid, pH 7.0 | **No. of used reflections** | 153739 (15024) |
| **Beamline** | Diamond I04 | Diamond I03 | **R factor (%)** | 17.7 |
| **Wavelength (Å)** | 0.97941 | 0.97626 | **R free (%)** | 20.1 |
| **Oscillation (º)** | 0.2 | 0.2 | **Most favourable Ramachandran region (%)** | 97.69 |
| **Total oscillation (º)** | 480 | 180 | **Generously allowed regions (%)** | 2.31 |
| **Unit cell parameters (Å, degree)** | a = 101.54, b = 123.95, c = 74.28 90, 104.89, 90 | a = 87.14, b = 140.81, c = 74.03 90, 98.65, 90 | **Disallowed regions (%)** | 0 |
| | | | **Bond distance (Å)** | 0.005 |
| | | | **Angles (º)** | 0.77 |
| **Space group** | C2 | C2 | | |
| **Resolution range (Å)** | 71.79-2.05 (2.10-2.05) | 55.50-1.45 (1.50-1.45) | | |
| **Total reflections** | 383069 (28104) | 293652 (26742) | | |
| **Unique reflections** | 55609 (4128) | 154410 (15025) | | |
| **R-merge** | 0.047 (1.182) | 0.041 (1.441) | | |
| **I/σ (I)** | 19.2 (1.5) | 9.93 (0.85) | | |
| **Completeness (%)** | 100.0 (100.0) | 99.4 (97.2) | | |
| **Multiplicity** | 6.9 (6.8) | 1.9 (1.8) | | |
| **Anomalous completeness (%)** | 99.6 (99.8) | | | |
| **Anomalous multiplicity** | 3.5 (3.4) | | | |
| **Anomalous correlation** | 0.320 (0.033) | | | |
| **Anomalous slope** | 1.68 | | | |

**Table S2. Data collection and refinement statistics for N-terminal Malonyl-CoA reductase (MCR-N) structure.** Values in brackets refer to high resolution shell.

| Data collection | Se-Met MCR-N | Native MCR-N | Refinement | Final MCR-N PDB: 8AEW |
|---|---|---|---|---|
| **Crystallisation condition** | 0.1 M Sodium cacodylate, pH 6.5, 40% (w/v) MPD 5% (w/v) PEG 8000 | 0.1 M Sodium HEPES, pH 7.5, 70% (w/v) MPD | **No. of used reflections** | 33849 (3195) |
| **Beamline** | Diamond I04 | Diamond I04-1 | **R factor (%)** | 24.5 |
| **Wavelength (Å)** | 0.97949 | 0.92819 | **R free (%)** | 29.2 |
| **Oscillation (º)** | 0.2 | 0.2 | **Most favourable Ramachandran region (%)** | 95.23 |
| **Total oscillation (º)** | 480 | 180 | **Generously allowed regions (%)** | 4.77 |
| **Unit cell parameters (Å, degree)** | a = 54.12, b = 84.75, c = 257.27 90, 90, 90 | a = 54.76, b = 86.79, c = 260.49 90, 90, 90 | **Disallowed regions (%)** | 0.00 |
| | | | **Bond distance (Å)** | 0.008 |
| | | | **Angles (º)** | 1.12 |
| **Space group** | $P2_12_12_1$ | $P2_12_12_1$ | | |
| **Resolution range (Å)** | 70.77-2.97 (3.05-2.97) | 82.34-2.72 (2.82-2.72) | | |
| **Total reflections** | 332602 (24475) | 68074 (6553) | | |
| **Unique reflections** | 25320 (1833) | 34118 (3195) | | |
| **R-merge** | 0.210 (1.965) | 0.109 (1.273) | | |
| **I/σ (I)** | 10.0 (1.2) | 7.5 (0.7) | | |
| **Completeness (%)** | 99.7 (99.3) | 98.5 (96.1) | | |
| **Multiplicity** | 13.1 (13.4) | 2.0 (2.0) | | |
| **Anomalous completeness (%)** | 99.8 (98.9) | | | |
| **Anomalous multiplicity** | 7.0 (7.1) | | | |
| **Anomalous correlation** | 0.354 (0.030) | | | |
| **Anomalous slope** | 1.163 | | | |

**Table S3. Data collection and refinement statistics for NADP-bound, and NADP and malonate-bound MCR-C structures.** Values in brackets refer to high resolution shell.

| | NADP-bound<br>PDB: 8A7S | NADP and malonate-bound<br>PDB: 8A8T |
|---|---|---|
| **Crystallisation condition** | 0.8 M succinic acid, pH 7.0 | |
| **Ligand** | 1 mM NADPH soaked | 0.5 mM NADPH co-crystallised |
| **Beamline** | Diamond I03 | Diamond I04 |
| **Wavelength (Å)** | 0.97626 | 0.97949 |
| **Unit cell parameters (Å, degree)** | a = 87.20, b = 140.72<br>c = 74.31<br>90, 98.41, 90 | a = 100.96, b = 126.29<br>c = 74.33<br>90, 104.84, 90 |
| **Space group** | C2 | C2 |
| **Resolution range (Å)** | 49.03-1.54 (1.60-1.54) | 63.15-2.11 (2.18-2.11) |
| **Total reflections** | 212003 (8033) | 96890 (10212) |
| **Unique reflections** | 114540 (4951) | 50822 (5221) |
| **R-merge** | 0.046 (0.449) | 0.038 (0.535) |
| **I/σ (I)** | 4.2 (0.56) | 11.1 (1.4) |
| **Completeness (%)** | 86.5 (38.2) | 98.2 (99.9) |
| **Multiplicity** | 1.9 (1.5) | 1.9 (2.0) |
| **No. of used reflections** | 112295 (4951) | 50801 (5221) |
| **R factor (%)** | 17.1 | 18.2 |
| **R free (%)** | 19.9 | 21.7 |
| **Most favourable Ramachandran region (%)** | 98.16 | 97.48 |
| **Generously allowed regions (%)** | 1.69 | 2.52 |
| **Disallowed regions (%)** | 0.15 | 0.00 |
| **Bond distance (Å)** | 0.006 | 0.007 |
| **Angles (°)** | 0.80 | 0.83 |

**Table S4. Structural alignment of all MCR-C structures.** RMSD values (Å) are given for all alignments. Calculated using GE[1] (General Efficient Structural Alignment of Macromolecular Targets) in CCP4.

| Structure | MCR_C Apo | NADPH-bound | NADPH and malonate-bound | R773Q | R773A | Y731A | E779W |
|---|---|---|---|---|---|---|---|
| MCR_C Apo | 0 | 0.291 | 1.379 | 1.184 | 0.297 | 1.258 | 0.256 |
| NADPH-bound | | 0 | 1.320 | 1.315 | 0.353 | 1.304 | 0.350 |
| NADPH and malonate-bound | | | 0 | 0.512 | 1.225 | 0.412 | 1.254 |
| R773Q | | | | 0 | 1.185 | 0.246 | 1.325 |
| R773A | | | | | 0 | 1.215 | 0.323 |
| Y731A | | | | | | 0 | 1.303 |
| E779W | | | | | | | 0 |

**Table S5. Crystallisation conditions for MCR-C**

| Condition | Screen-No. | Structures |
|---|---|---|
| 0.8 M Succinic acid pH 7.0 | Molecular Dimensions JCSG+ 2-19 | Native MCR-C, NADPH-bound MCR-C, NADPH/malonate-bound MCR-C |
| 0.1 M Sodium cacodylate trihydrate pH 6.5, 1.4 M Sodium acetate trihydrate | Hampton Research Screen 1-7 | Se-Met MCR-C, R773Q MCR-C, E779W MCR-C |
| 0.1 M Sodium citrate tribasic dihydrate, 1 M Ammonium phosphate | Hampton Research Screen 1-11 | R773A MCR-C, Y731A MCR-C |
| 0.2 M Lithium sulfate, 0.1 M phosphate/citrate, 20 % PEG 1000 | Molecular Dimensions JCSG+ 1-6 | |
| 1 M Ammonium sulfate, 0.1 M Bis Tris, 1 % PEG 3350 | Molecular Dimensions JCSG+ 2-38 | |
| 0.1 M Sodium HEPES pH 7.5, 0.8 M Sodium potassium tartrate tetrahydrate | Hampton Research Screen 1-29 | |

**Table S6. Crystallisation conditions for MCR-N**

| Condition | Screen-No. | Structures |
|---|---|---|
| 0.1 M Na cacodylate, 40% MPD 5% PEG 8000 | Molecular Dimensions JCSG+ 1-17 | Se-Met MCR-N |
| 0.1 M Na HEPES, pH 7.5, 70% MPD | Molecular Dimensions JCSG+ 1-41 | Native MCR-N |
| 0.2 M $MgCl_2$, 0.1 M Tris, 20% PEG 8000 | Molecular Dimensions JCSG+ 1-42 | |

**Table S7. Data collection and refinement statistics for MCR-C variant structures.**

| MCR-C variant | R773A PDB: 8AEO | Y731A PDB: 8AER | R773Q PDB: 8AEQ | E779W PDB: 8AET |
|---|---|---|---|---|
| **Crystallisation condition** | 0.1 M Sodium citrate tribasic dihydrate, 1 M Ammonium phosphate | | 0.1 M Sodium cacodylate, pH 6.5, 1.4 M Sodium acetate trihydrate | |
| **Beamline** | Diamond I03 | Diamond I03 | Diamond I03 | Diamond I03 |
| **Wavelength (Å)** | 0.97625 | 0.97625 | 0.97625 | 0.97625 |
| **Temperature (K)** | 100 | 100 | 100 | 100 |
| **Detector** | Pilatus3 6M | Pilatus3 6M | Pilatus3 6M | Pilatus3 6M |
| **Oscillation (º)** | 0.2 | 0.2 | 0.2 | 0.2 |
| **Total oscillation (º)** | 180 | 180 | 180 | 180 |
| **Space group** | C2 | C2 | C2 | C2 |
| **Unit cell parameters (Å, º)** | a = 87.71<br>b = 140.26<br>c = 73.84<br>90, 98.84, 90 | a = 101.71<br>b = 124.51<br>c = 74.68<br>90, 105.09, 90 | a = 102.07<br>b = 123.65<br>c = 74.25<br>90, 105.50, 90 | a = 87.17<br>b = 139.64<br>c = 73.31<br>90, 98.29, 90 |
| **Resolution range (Å)** | 41.15-1.76 (1.83-1.76) | 48.00-1.77 (1.83-1.77) | 46.78-1.64 (1.69-1.64) | 48.69-2.00 (2.07-2.00) |
| **Total reflections** | 161792 (16583) | 156726 (15751) | 196275 (19284) | 112286 (10828) |
| **Unique reflections** | 85187 (8578) | 83838 (8440) | 104921 (10207) | 58178 (5614) |
| **R-merge** | 0.034 (0.354) | 0.044 (0.399) | 0.028 (0.834) | 0.055 (0.429) |
| **I/σ (I)** | 10.6 (1.9) | 9.4 (1.5) | 11.1 (1.6) | 7.4 (1.5) |
| **CC half** | 0.9 (0.6) | 1.0 (0.6) | 1.0 (0.4) | 1.0 (0.7) |
| **Completeness (%)** | 98.3 (99.4) | 96.2 (97.1) | 96.6 (94.6) | 97.9 (96.7) |
| **Multiplicity** | 1.9 (1.9) | 1.9 (1.9) | 1.9 (1.9) | 1.9 (1.9) |
| **No. of used reflections** | 85178 (8578) | 83837 (8440) | 104613 (10207) | 57210 (5615) |
| **R factor (%)** | 17.3 | 17.4 | 17.9 | 19.1 |
| **R free (%)** | 20.3 | 19.7 | 20.3 | 22.1 |
| **Most favourable Ramachandran region (%)** | 98.39 | 97.90 | 98.05 | 98.75 |
| **Generously allowed regions (%)** | 1.61 | 1.94 | 1.79 | 1.25 |
| **Disallowed regions (%)** | 0.00 | 0.16 | 0.16 | 0.00 |
| **Bond distance (Å)** | 0.006 | 0.007 | 0.006 | 0.006 |
| **Angles (º)** | 0.76 | 1.01 | 0.78 | 0.82 |

**Table S8. Primers used for site-directed mutagenesis of MCR.** The nucleotide changes for mutations are highlighted in bold. The melting temperatures of primers were calculated with NEBaseChanger®.

| Mutation | Primer | Sequence (5' to 3') | Length (bp) | $T_m$ (ºC) |
|---|---|---|---|---|
| R734A | Forward | CTACCCCAAC**GC**TGCCGATTACGCC | 25 | 59 |
| | Reverse | GGAATGGCCGCATCTTTTTC | 20 | 62 |
| R734Q | Forward | TACCCCAACC**AG**GCCGATTACG | 22 | 59 |
| | Reverse | GGGAATGGCCGCATCTTT | 18 | 63 |
| Y731A | Forward | GGCCATTCCC**GCA**CCCAACCGTGC | 24 | 58 |
| | Reverse | GCATCTTTTTCACCGCCAAAG | 21 | 63 |
| R773A | Forward | CGAAGGTGAT**GC**CTTGCGCGGTACCG | 26 | 67 |
| | Reverse | ACCGGACCCGGCGCAATG | 18 | 72 |
| R773Q | Forward | GAAGGTGATC**AG**TTGCGCGGTACCGGTGAACG | 32 | 68 |
| | Reverse | GACCGGACCCGGCGCAAT | 18 | 72 |
| E779W | Forward | CGGTACCGGT**TGG**CGTCCCGGCC | 23 | 68 |
| | Reverse | CGCAAGCGATCACCTTCG | 18 | 64 |

**Table S9. Activity determination of MCR-C.** Initial rates were calculated for five replicates of 0.1 mM malonyl-CoA, and converted to activities.

| No. | Malonyl-CoA (mM) | Rate (AU $sec^{-1}$) | Activity (µmol $min^{-1}$ $mg^{-1}$ protein) |
|---|---|---|---|
| 1 | 0.1 | 0.0041 | 1.952 |
| 2 | 0.1 | 0.0037 | 1.762 |
| 3 | 0.1 | 0.0044 | 2.095 |
| 4 | 0.1 | 0.0032 | 1.524 |
| 5 | 0.1 | 0.0040 | 1.905 |
| Average activity: 1.85 ± 0.22 µmol $min^{-1}$ $mg^{-1}$ protein (n=5) | | | |

**Table S10. Activity determination of MCR-N.** Initial rates were calculated for five replicates of products of MCR-C reactions with 0.1 mM malonyl-CoA, and they were converted to the activity values.

| No. | Malonyl-CoA (mM) | Rate (AU $sec^{-1}$) | Activity (μmol $min^{-1}$ $mg^{-1}$ protein) |
|---|---|---|---|
| 1 | 0.1 | 0.0009 | 0.426 |
| 2 | 0.1 | 0.0008 | 0.381 |
| 3 | 0.1 | 0.0008 | 0.381 |
| 4 | 0.1 | 0.0007 | 0.333 |
| 5 | 0.1 | 0.0007 | 0.333 |
| Average activity: 0.37 ± 0.04 μmol $min^{-1}$ $mg^{-1}$ protein (n=5) | | | |

**Table S11. Determination of Michaelis-Menten parameters of C-terminal MCR.** Initial rates were calculated for various malonyl-CoA concentrations. Michaelis-Menten parameters were determined using average rates by Lineweaver-Burk plot method.

| Malonyl-CoA (mM) | Rate (AU $sec^{-1}$) | Rate (μmol $min^{-1}$) |
|---|---|---|
| 0.05 | 0.00330±0.00028 | 0.0157±0.0013 (n=2) |
| 0.075 | 0.00370±0.00028 | 0.0176±0.0013 (n=2) |
| 0.1 | 0.00388±0.00041 | 0.0185±0.0019 (n=5) |
| 0.125 | 0.00392±0.00073 | 0.0187±0.0034 (n=5) |
| 0.15 | 0.00397±0.00075 | 0.0189±0.0035 (n=3) |
| $K_m$= 0.0174 ± 0.0030 mM, $V_{max}$= 0.0214 ± 0.0037 μmol $min^{-1}$ | | |

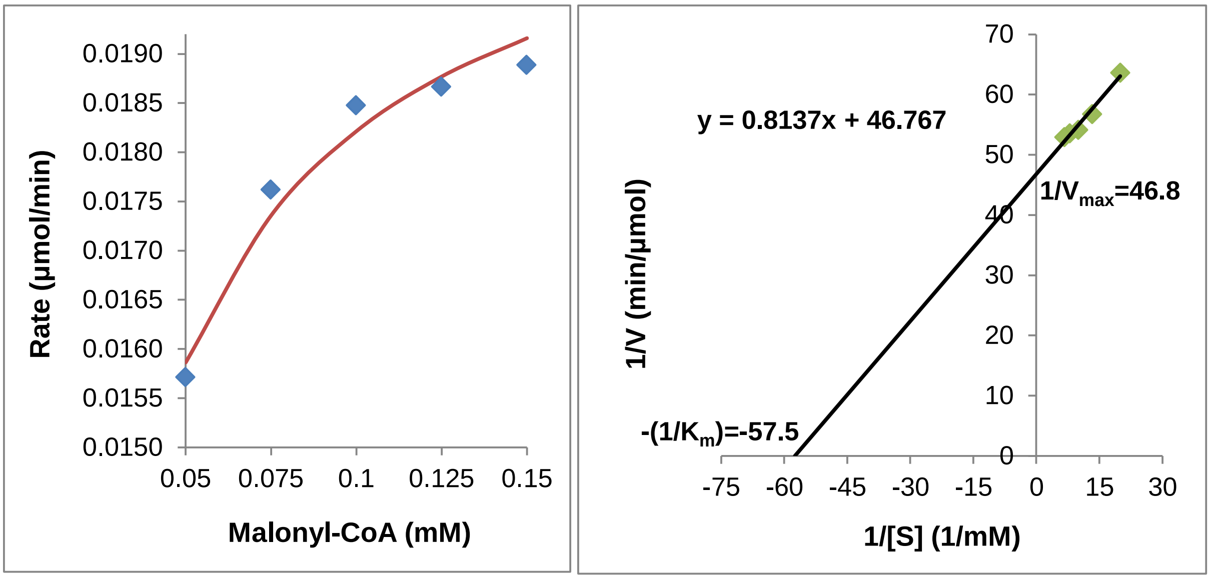


**Figure S1. Determination of Michaelis-Menten parameters of C-terminal MCR.** Rate vs. Malonyl-CoA plot (left), and Lineweaver-Burk plot (right) using average rates. According to the Lineweaver-Burk plot (y=0.8137x + 46.767), $1/V_{max}$, and $K_m/V_{max}$ was 46.767 min $\mu mol^{-1}$ and

0.8137 mM min μmol$^{-1}$, respectively.

**Table S12. Determination of Michaelis-Menten parameters of N-terminal MCR.** Initial rates were calculated for various initial malonyl-CoA concentrations. Michaelis-Menten parameters were determined using average rates by Lineweaver-Burk plot method.

| Malonyl-CoA (mM) | Rate (AU sec$^{-1}$) | Activity (μmol min$^{-1}$) |
|---|---|---|
| 0.05 | 0.00039±1.41×10$^{-5}$ | 0.0019±0.00007 (n=2) |
| 0.075 | 0.00057±5.75×10$^{-5}$ | 0.0027±0.00027 (n=3) |
| 0.1 | 0.00078±7.48×10$^{-5}$ | 0.0037±0.00036 (n=5) |
| 0.125 | 0.00097±5.77×10$^{-5}$ | 0.0046±0.00022 (n=3) |
| 0.15 | 0.00102±7.64×10$^{-5}$ | 0.0048±0.00036 (n=3) |
| $K_m$= 1.27 ± 0.23 mM, $V_{max}$= 0.049 ± 0.009 μmol min$^{-1}$ | | |

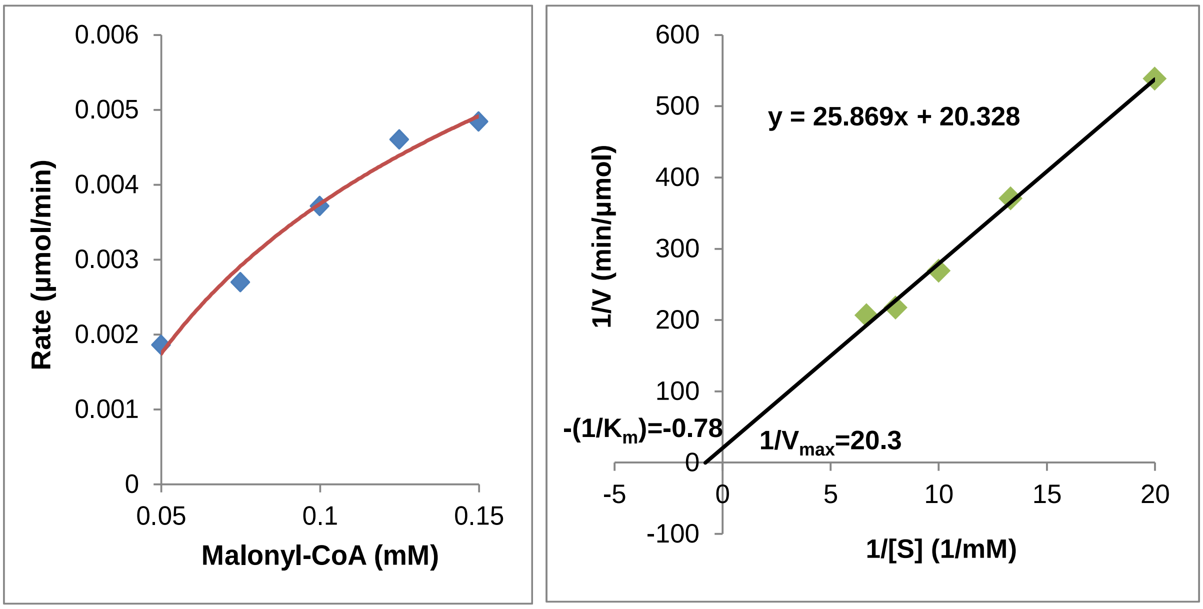


**Figure S2. Determination of Michaelis-Menten parameters of N-terminal MCR.** NADPH oxidation was monitored at 340 nm for various malonyl-CoA concentrations (0.05, 0.075, 0.1, 0.125, and 0.15 mM) (Absorption vs. Time plots). Reaction rates were determined based on initial rates for each concentration, and Rate vs. Malonyl-CoA plot was generated using average rates.

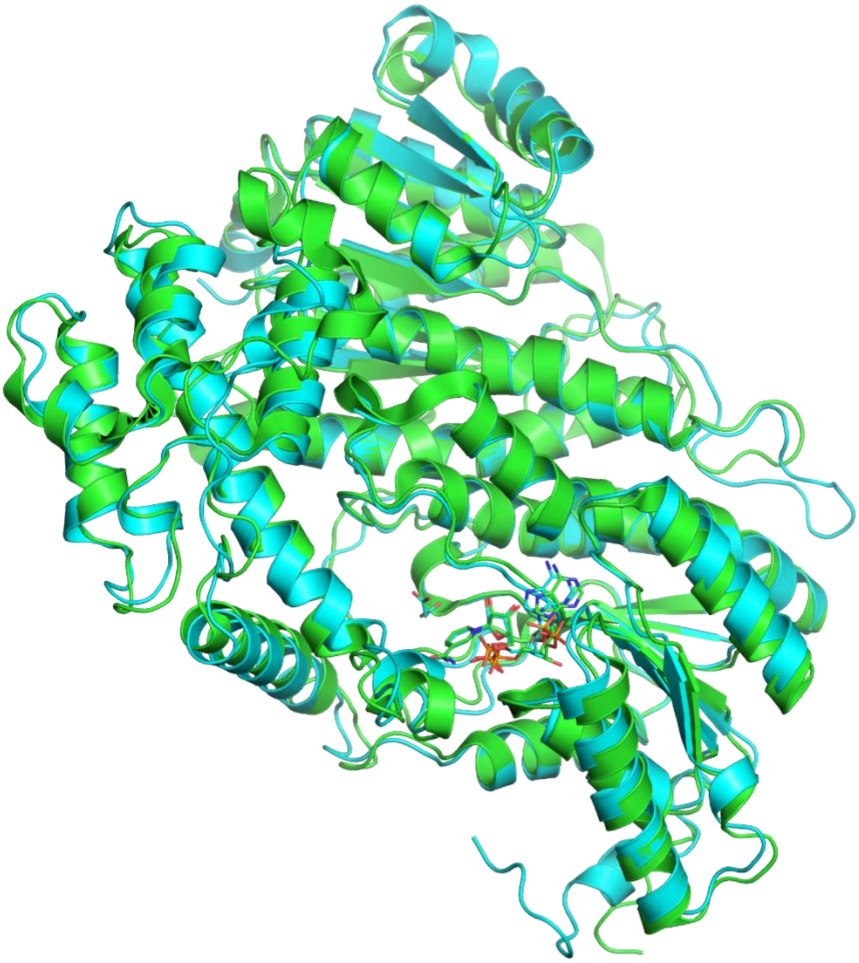

**Figure S3.** Superposition of NADP-bound MCR-C structures from *C. aurantiacus* (turquoise) and *P. dokdonensis* (green)[2] (PDB ID: 6K8U) (RMSD=0.9 Å). NADPs and malonate are shown as sticks. NADPs are found at the same site.

A

B

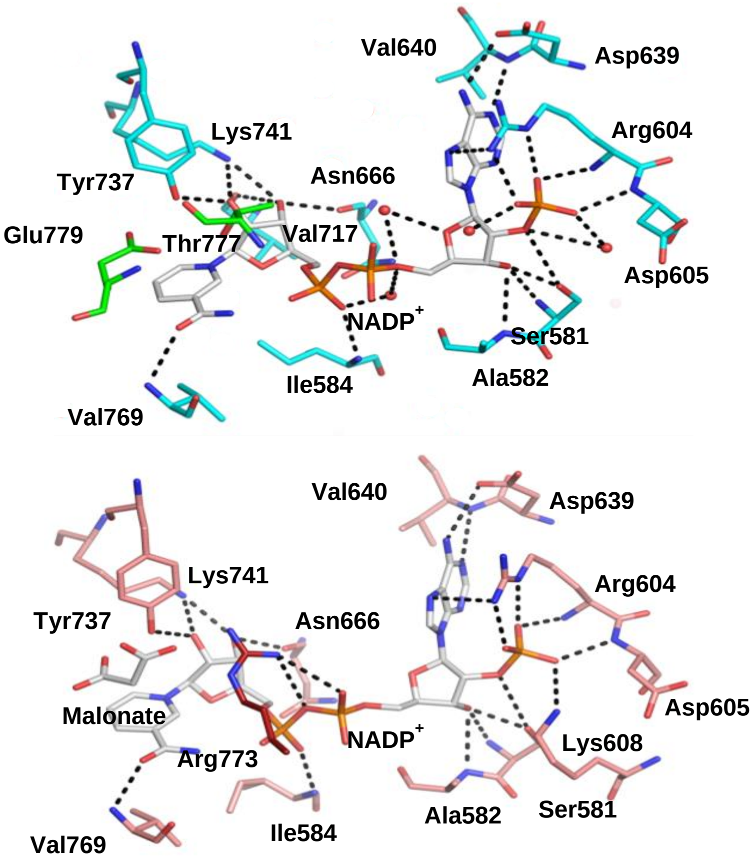


**Figure S4. Interactions of $NADP^+$ in NADP-bound (A), and NADP/malonate-bound (B) C terminal MCR structures.** The residues are shown in turquoise sticks, loop residues are shown in green, in the NADP-bound structure. The residues are shown in salmon sticks, Arg773 in the loop is shown in red, in the NADP/malonate-bound structure. $NADP^+$ and malonate are shown in white sticks. Water molecules are represented with red spheres.

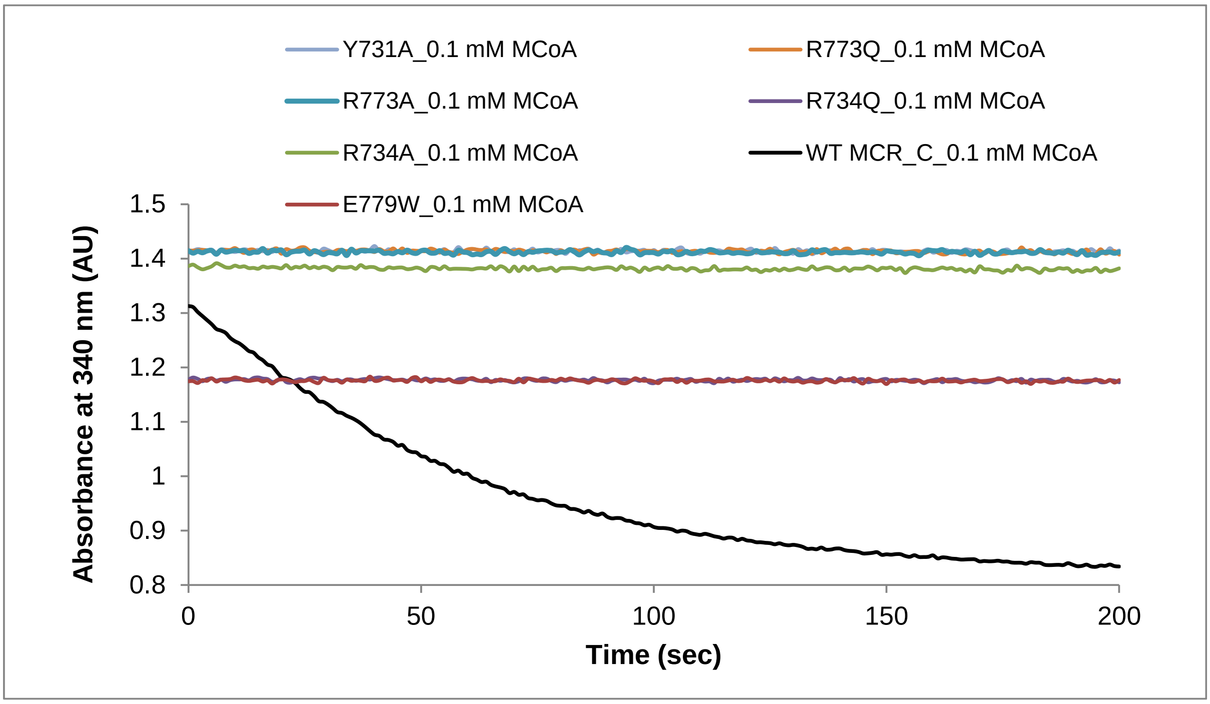


**Figure S5. Enzyme assays of MCR variants.** NADPH oxidation was monitored at 340 nm using 0.1 mM malonyl-CoA (MCoA) for MCR variants; Y731A, R773A, R734A, E779W, R773Q, and R734Q, No NADPH oxidation was obtained in contrast with wild type (WT) MCR-C.

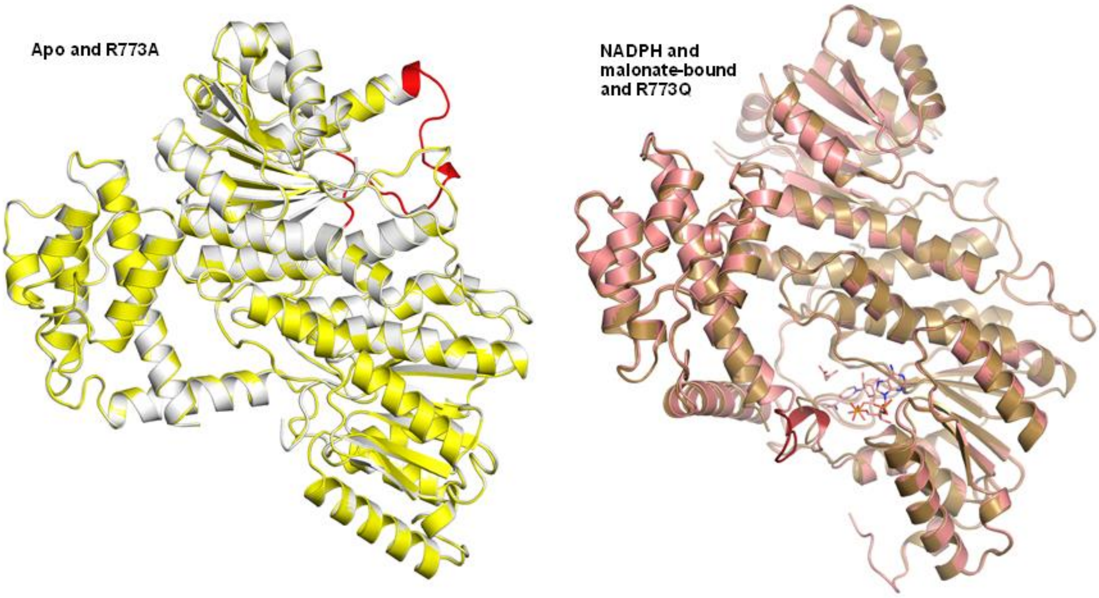


**Figure S6. Superposition of apo and R773A structures, and NADPH and malonate-bound and R773Q MCR-C structures.** Apo and R773A (left), and NADPH and malonate-bound and R773Q (right) structures are shown in cartoon view, white, yellow, salmon, and brown respectively. The disordered regions in R773A and R773Q are represented in apo and NADPH and malonate-bound structures in red. The disordered regions in R773A are also disordered in both NADPH and malonate-bound, and R773Q structures. R773A is in “open” conformation while R773Q is in “closed” conformation. NADPH and malonate are shown as sticks in salmon (right).

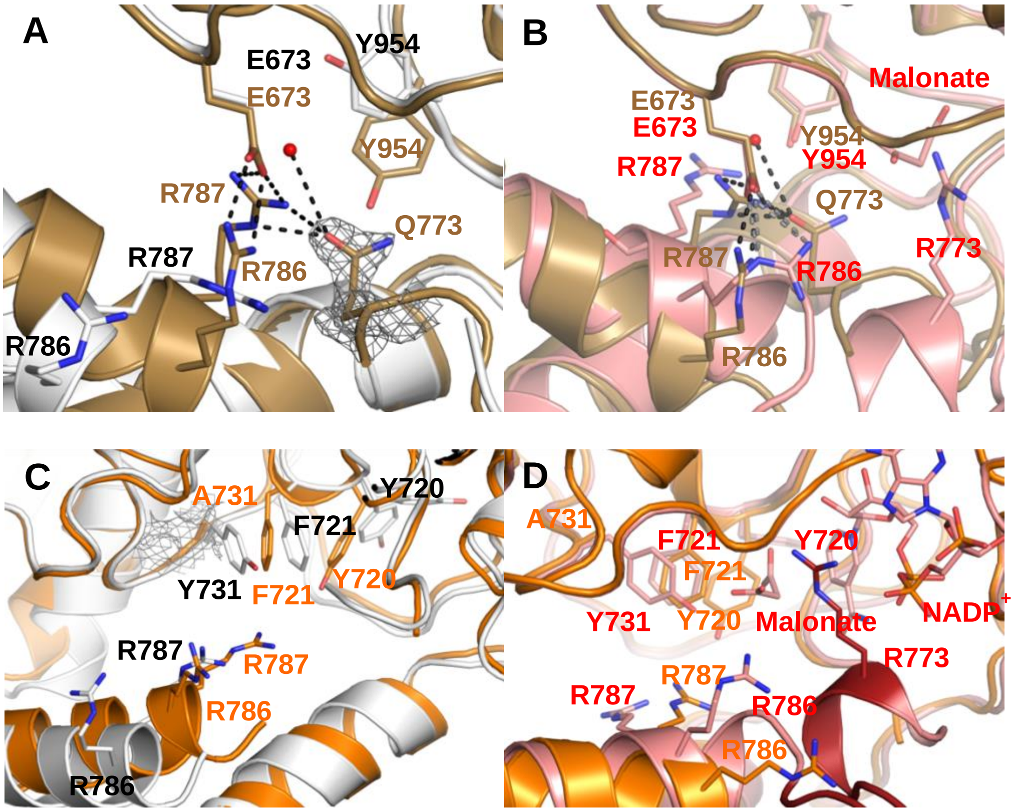


**Figure S7. A-B**, **Comparison of R773Q interactions with apo and NADPH/malonate-bound MCR-C structures.** R773Q superposed with apo (A), and NADPH/malonate bound (B) structures, shown in cartoon view, brown, white, and salmon respectively. The interacting residues and malonate are shown as sticks, and labelled brown, black, and red for R773Q, apo, and NADPH/malonate bound structures, respectively. Hydrogen bonds are shown in black and grey in R773Q and NADPH/malonate bound structures, respectively. **C-D, Comparison of Y731A interactions with apo and NADPH/malonate-bound MCR-C structures.** Y731A superposed with apo (C), and NADPH/malonate bound (D) structures, shown in cartoon view, orange, white, and salmon respectively. The interacting residues, malonate, and NADPH are shown as sticks, and labelled orange, black, and red for Y731A, apo, and NADPH/malonate bound structures, respectively. Electron density map (2mFo-2DFc) contoured at 1.3 sigma around Gln773 and Ala731 are shown in grey mesh (A and C). Water molecule is shown as red sphere (A and B).

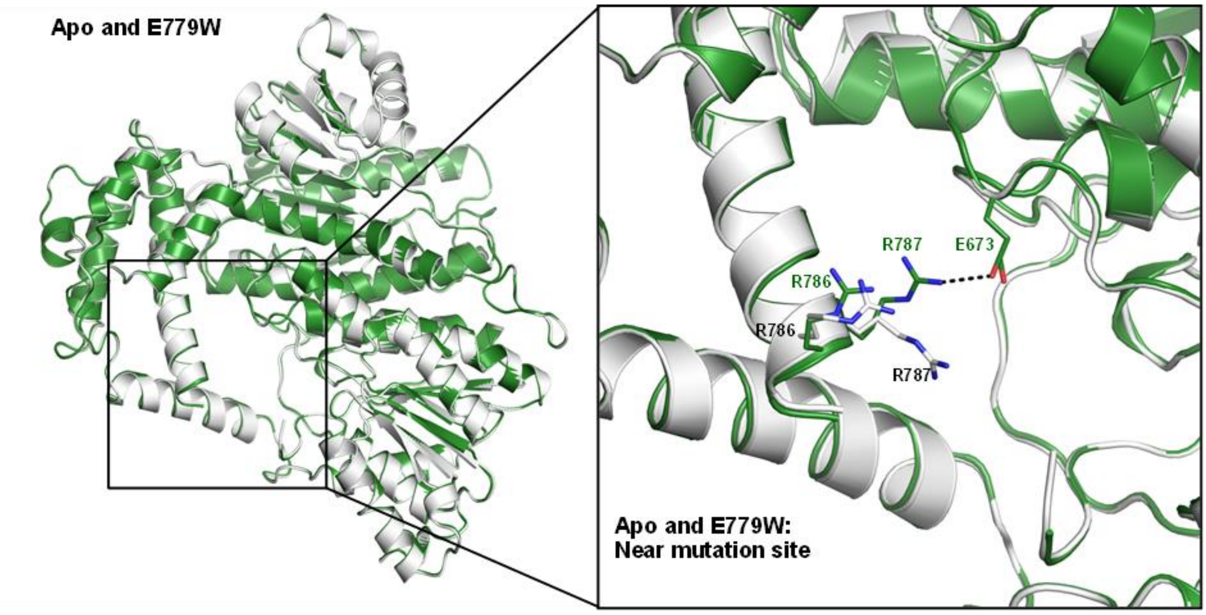


**Figure S8. Superposition of E779W, and apo MCR-C structures.** Apo and E779W structures are shown in cartoon view, white, and green, respectively. E779W is in “open” conformation. Conformational changes near mutation site are shown in the right. Residues are represented as sticks, and labelled black and green for apo, and E779W structures, respectively.

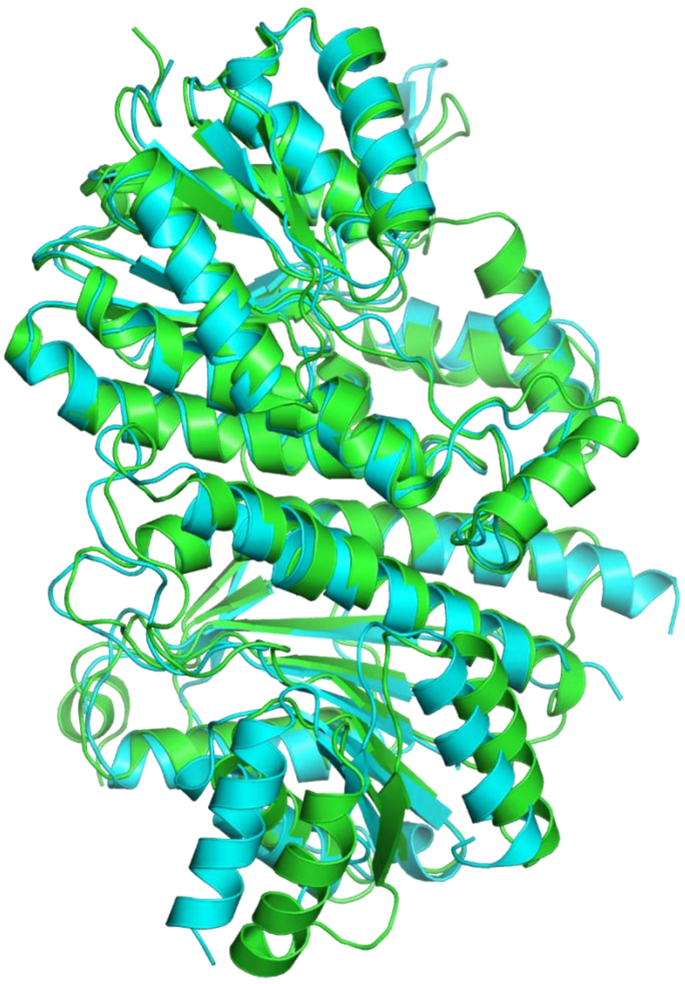

**Figure S9.** Superposition of MCR-N structures from *C. aurantiacus* (turquoise) and *P. dokdonensis* (green)[2] (PDB ID: 6K8V) (RMSD=1.2 Å).

| | | |
|---|---|---|
| malonyl-CoA reductase [*Chloroflexus aurantiacus*] | $_{15}$TGGAGNIG$_{22}$ .. | $_{155}$NVSTIFSRAEYYGRIPYVTPK$_{175}$ .. |
| malonyl-CoA reductase [*Porphyrobacter dokdonensis* DSW-74] | $_{29}$TGAAGNLG$_{36}$ .. | $_{175}$NVSTIFSRTPYYARAAYVVPK$_{195}$ .. |
| NAD(P)-dependent oxidoreductase [*Chloroflexus sp.* MS-G] | $_{15}$TGGAGNIG$_{22}$ .. | $_{155}$NVSTIFSYAEYYGRIPYVTPK$_{175}$ .. |
| NAD(P)-dependent oxidoreductase [*Chloroflexus sp.* Y-396-1] | $_{15}$TGGAGNIG$_{22}$ .. | $_{155}$NISTIFSYAEYYGRIPYVTPK$_{175}$ .. |
| short-chain dehydrogenase [*Chloroflexus aggregans*] | $_{15}$TGGAGNIG$_{22}$ .. | $_{155}$NISTIFSRAEYYGRIPYVVPK$_{175}$ .. |
| short-chain dehydrogenase [*Chloroflexus islandicus*] | $_{15}$TGGAGNIG$_{22}$ .. | $_{155}$NISTIFSRAEYYGRIPYVAPK$_{175}$ .. |
| short-chain dehydrogenase [*Oscillochloris trichoides*] | $_{14}$TGGAGTIG$_{21}$ .. | $_{154}$NISTIFARTDYYGRIPYVVPK$_{174}$ .. |
| short-chain dehydrogenase [*Roseiflexus castenholzii*] | $_{15}$TGGAGNIG$_{22}$ .. | $_{155}$NISTIFSRTDYYGRIAYVAPK$_{175}$ .. |
| short-chain dehydrogenase [*Roseiflexus sp.* RS-1] | $_{18}$TGGAGNIG$_{25}$ .. | $_{158}$NVSTIFSRTDYYGRIAYVVPK$_{178}$ .. |
| hypothetical protein HY22_11770 [*Chlorobium sp.* GBChlB] | $_{5}$TGGAGNIG$_{12}$ .. | $_{145}$NVSTIFSRTEYFGRAAYSVPK$_{165}$ .. |
| NAD-dependent dehydratase [*Erythrobacter sp.* SCN 62-14] | $_{29}$TGAAGNLG$_{36}$ .. | $_{175}$NVSTIFSRTPYYARAAYVVPK$_{195}$ .. |
| NAD-dependent dehydratase [*Methylibium sp.* NZG] | $_{15}$TGAAGNLG$_{22}$ .. | $_{155}$NVSTIFSRTNYYGRTAYVVPK$_{175}$ .. |
| NAD-dependent dehydratase [*Porphyrobacter sp.* AAP60] | $_{24}$TGAAGNLG$_{31}$ .. | $_{164}$NVSTIFSRTPYYARAAYVVPK$_{184}$ .. |
| NAD-dependent dehydratase [*Proteobacteria*] | $_{31}$TGAAGNLG$_{38}$ .. | $_{166}$NISTIFSHTRYYGRTAYVVPK$_{186}$ .. |
| NAD-dependent dehydratase [*Alphaproteobacteria*] | $_{29}$TGAAGNLG$_{36}$ .. | $_{169}$NVSTIFSRTPYYARAAYVVPK$_{189}$ .. |
| short-chain dehydrogenase [*gamma proteobacterium* NOR5-3] | $_{25}$TGAAGNIG$_{32}$ .. | $_{165}$NVSTIFSRTHYYGRIPYVVPK$_{185}$ .. |
| NAD-dependent epimerase [*Erythrobacter sp.* NAP1] | $_{20}$TGAAGNLG$_{27}$ .. | $_{160}$NISTIFSHTRYYGRTAYVVPK$_{180}$ .. |
| NAD-dependent dehydratase [*Erythrobacter litoralis*] | $_{21}$TGAAGNLG$_{28}$ .. | $_{161}$NISTIFSHTRYYGRTAYVVPK$_{181}$ .. |
| NAD-dependent dehydratase [*Novosphingobium sp.* AAP83] | $_{15}$TGAAGNLG$_{22}$ .. | $_{155}$NVSTIFSRTPYYARAAYVVPK$_{175}$ .. |

**Figure S10. Sequence alignment of the N-terminal MCR with SDR proteins.** Regions for dinucleotide binding motif (TGXXX[AG]XG), active site motif (YXXXK), and putative catalytic residues are shown. Conserved and non-conserved residues are shown in red, and black, respectively. YXXXK regions are shown in blue. Putative catalytically important residues [Tyr and Arg (Tyr165 and Arg168 in the MCR-N)] are shown in green. Glu164 (coloured in purple) is unlikely a catalytic residue. The alignment was carried out using BLAST.

| | | | | | |
|---|---|---|---|---|---|
| malonyl-CoA reductase [*Chloroflexus aurantiacus*] | $_{578}$TGGSAGIG$_{585}$ | .. | $_{719}$SYFGGEKDAAIPYPNRADYAVSK$_{741}$ | .. | $_{771}$GDRLRGTGE$_{779}$ |
| malonyl-CoA reductase [*Porphyrobacter dokdonensis* DSW-74] | $_{585}$TGGSAGIG$_{592}$ | .. | $_{726}$SYFGGEKYLAVAYPNRADYAVSK$_{748}$ | .. | $_{778}$GDRLSGTGG$_{789}$ |
| short-chain dehydrogenase [*Chloroflexus sp.* MS-G] | $_{578}$TGGSAGIG$_{585}$ | .. | $_{719}$SYFGGEKEAAIPYPNRADYAVSK$_{741}$ | .. | $_{771}$GDRLRGTGE$_{779}$ |
| short-chain dehydrogenase [*Chloroflexus sp.* Y-396-1] | $_{578}$TGGSAGIG$_{585}$ | .. | $_{719}$SYFGGEKEAAIPYPNRADYAVSK$_{741}$ | .. | $_{771}$GDRLRGTGE$_{779}$ |
| short-chain dehydrogenase [*Chloroflexus aggregans*] | $_{578}$TGGSAGIG$_{585}$ | .. | $_{719}$SYFGGEKDAAIPYPNRADYAVSK$_{741}$ | .. | $_{771}$GDRLRGTGE$_{779}$ |
| short-chain dehydrogenase [*Chloroflexus islandicus*] | $_{578}$TGGSAGIG$_{585}$ | .. | $_{719}$SYFGGEKDAAIPYPNRSDYAVSK$_{741}$ | .. | $_{771}$GDRLRGTGE$_{779}$ |
| short-chain dehydrogenase [*Oscillochloris trichoides*] | $_{579}$TGGSAGIG$_{586}$ | .. | $_{720}$SYFGGEKYAAIPYPNRADYAVSK$_{742}$ | .. | $_{772}$GERLRGSGD$_{780}$ |
| short-chain dehydrogenase [*Roseiflexus castenholzii*] | $_{590}$TGGSAGIG$_{597}$ | .. | $_{731}$SYFGGEKYVAIPYPNRSDYAVSK$_{753}$ | .. | $_{783}$GERLKGAGS$_{791}$ |
| short-chain dehydrogenase [*Roseiflexus sp.* RS-1] | $_{593}$TGGSAGIG$_{600}$ | .. | $_{734}$SYFGGEKYVAIPYPNRSDYAVSK$_{756}$ | .. | $_{786}$GERLKGAAN$_{794}$ |
| hypothetical protein HY22_11770 [*Chlorobium sp.* GBChlB] | $_{551}$TGGSQGIG$_{558}$ | .. | $_{692}$SYFGGEKYVAVPYPNRADYAVSK$_{714}$ | .. | $_{744}$GIRLAGEGN$_{752}$ |
| NAD-dependent dehydratase [*Erythrobacter sp.* SCN 62-14] | $_{585}$TGGSAGIG$_{592}$ | .. | $_{727}$SYFGGEKYLAVAYPNRADYAVSK$_{749}$ | .. | $_{778}$GDRLSGTGG$_{787}$ |
| NAD-dependent dehydratase [*Methylibium sp.* NZG] | $_{569}$TGGSAGIG$_{576}$ | .. | $_{711}$SYFGGEKNLAVAYPNRSDYAVSK$_{732}$ | .. | $_{762}$GERLKGTGN$_{770}$ |
| NAD-dependent dehydratase [*Porphyrobacter sp.* AAP60] | $_{580}$TGGSAGIG$_{587}$ | .. | $_{721}$SYFGGEKYLAVAYPNRADYAVSK$_{743}$ | .. | $_{773}$GDRLSGTGG$_{781}$ |
| NAD-dependent dehydratase [*Proteobacteria*] | $_{582}$TGGSAGIG$_{589}$ | .. | $_{723}$SYFGGEKFLAVAYPNRADYAVSK$_{745}$ | .. | $_{775}$GDRLAGTGG$_{783}$ |
| NAD-dependent dehydratase [*Alphaproteobacteria*] | $_{585}$TGGSAGIG$_{592}$ | .. | $_{727}$SYFGGEKYLAVAYPNRADYAVSK$_{749}$ | .. | $_{778}$GDRLSGTGG$_{787}$ |
| short-chain dehydrogenase [*gamma proteobacterium* NOR5-3] | $_{575}$TGGSLGIG$_{582}$ | .. | $_{716}$SYFGGEKYVAVAYPNRADYAVSK$_{738}$ | .. | $_{768}$GARLRGLGG$_{776}$ |
| NAD-dependent epimerase [*Erythrobacter sp.* NAP1] | $_{572}$TGGSAGIG$_{579}$ | .. | $_{713}$SYFGGEKFLAVAYPNRADYGLSK$_{735}$ | .. | $_{765}$GDRLRGTGE$_{773}$ |
| NAD-dependent dehydratase [*Erythrobacter litoralis*] | $_{573}$TGGSAGIG$_{580}$ | .. | $_{714}$SYFGGEKFLAVAYPNRADYGLSK$_{736}$ | .. | $_{766}$GDRLSGTGG$_{774}$ |
| NAD-dependent dehydratase [*Novosphingobium sp.* AAP83] | $_{571}$TGGSAGIG$_{578}$ | .. | $_{712}$SYFGGEKYLAVAYPNRADYAASK$_{734}$ | .. | $_{764}$GDRLAGTGG$_{772}$ |

**Figure S11. Sequence alignment of the C-terminal MCR with SDR proteins.** Regions for dinucleotide binding motif (TGXXX[AG]XG), active site motif (YXXXK), and mutation sites are shown. Conserved and non-conserved residues are shown in red, and black, respectively. YXXXK regions are shown in brown. Catalytic Tyr (Tyr737 in the MCR-C), and other catalytically important residues [Ser, Tyr, Arg (Ser719, Tyr731, Arg734, Arg773 in the MCR-C)] are shown in purple, and green, respectively. The alignment was carried out using BLAST.

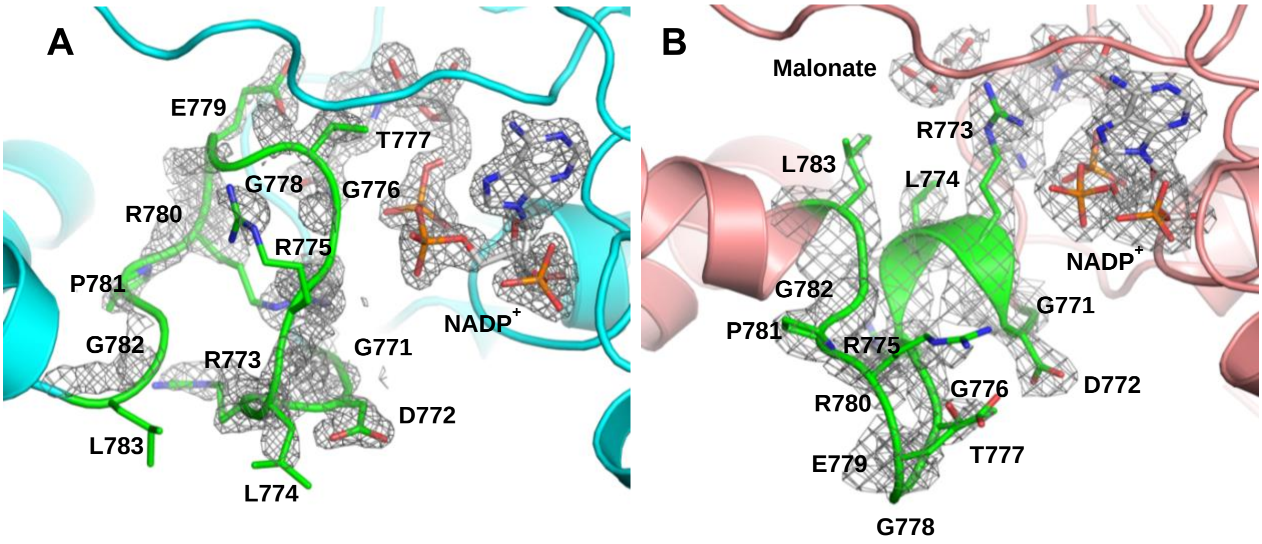


**Figure S12. The ordered loop in the NADP-bound (A) and NADP/malonate (B) MCR-C structures.** NADPH-bound and NADPH/malonate-bound structures are shown in cartoon view, turquoise and salmon, respectively. Electron density map ($2mF_o$-$2DF_c$) contoured at 1.3 sigma around the loop residues and $NADP^+$ is shown in grey mesh. The side chains in the loop are represented in green sticks.

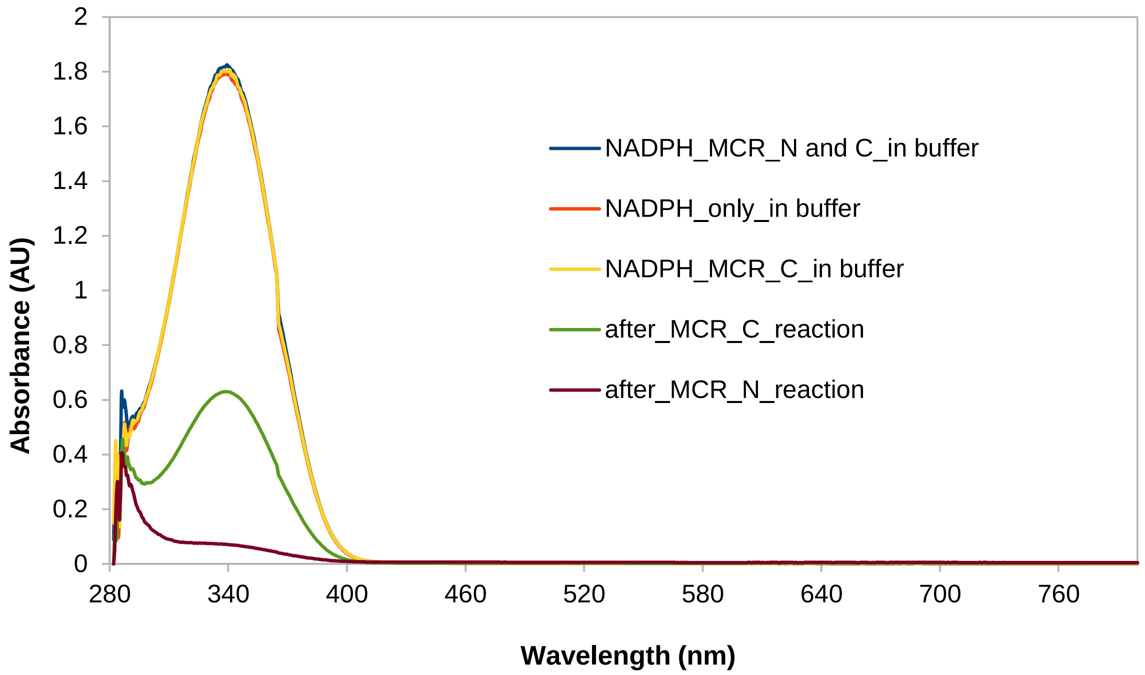


**Figure S13. Changes in spectra after the MCR reaction.** The spectra of the reaction medium were measured for; NADPH only in buffer, NADPH with MCR-C, NADPH with both enzymes (MCR-N and C), after the MCR-C reaction, after the MCR-N reaction. During the reaction, there was decrease in absorption at 340 nm. 0.2 mM malonyl CoA and 0.5 mM NADPH were used.

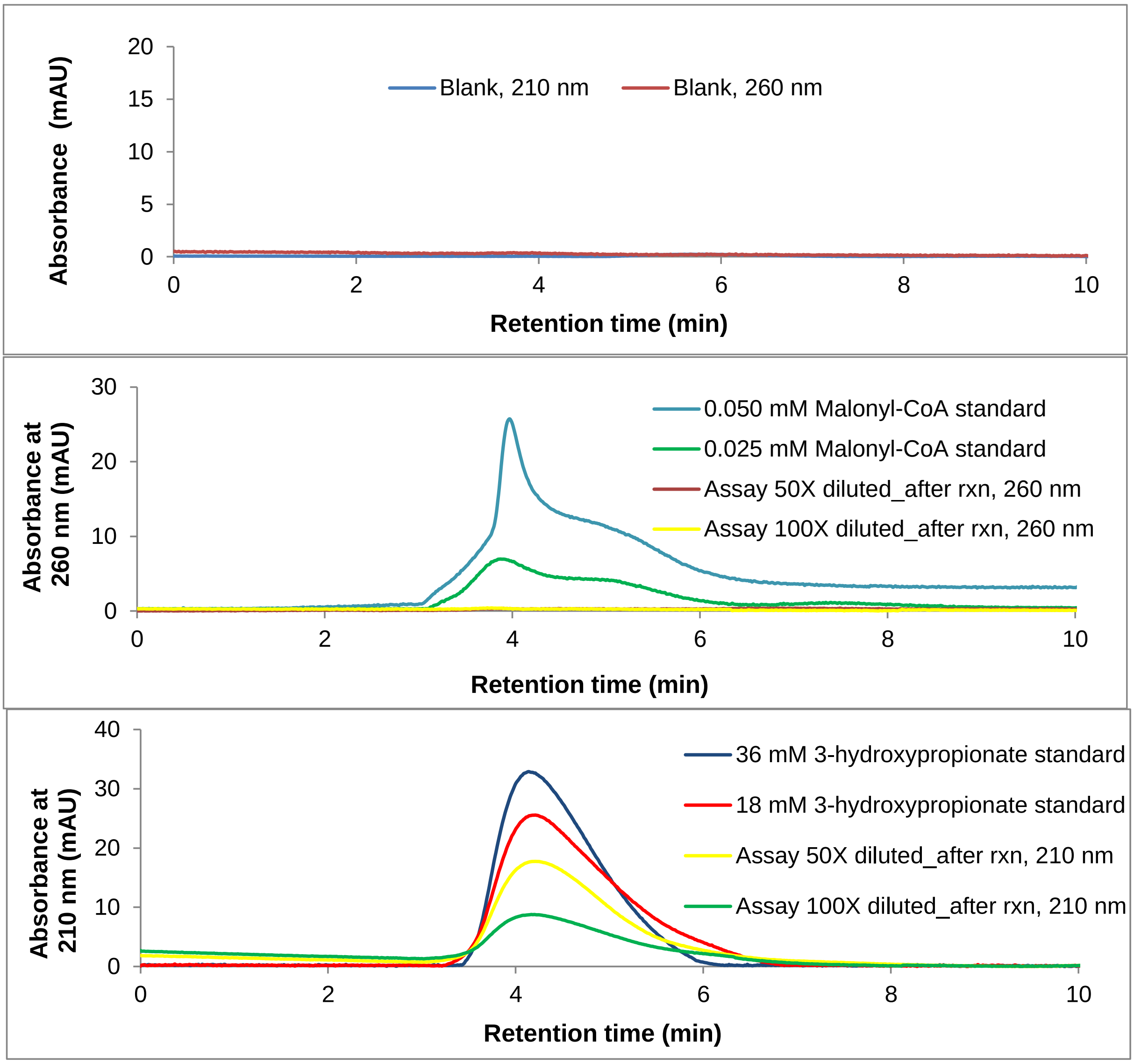


**Figure S14. HPLC analysis of MCR reaction product.** Absorbances at either 260 nm or 210 nm were measured for blank, malonyl CoA standard, 3 hydroxypropionate standard, and the assay reaction product. Mobile phase was used as a blank sample [40 mM $KH_2PO_4$, 0.2% formic acid, pH 4.2 : 70% methanol in 40 mM $KH_2PO_4$, 0.2% formic acid, pH 4.2 (50:50)]. Standards were prepared in mobile phase, and the reaction product was diluted with mobile phase (50-fold and 100-fold). Malonyl CoA (0.025 and 0.050 mM) was monitored at 260 nm, and 3 hydroxypropionate (18 and 36 mM) was monitored at 210 nm. Reaction product was monitored at both wavelengths.

**A**

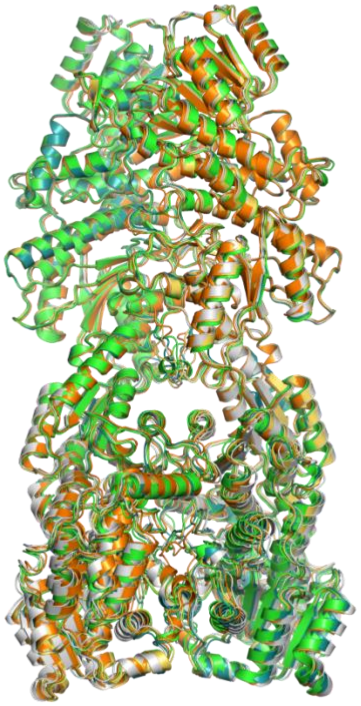

**B**

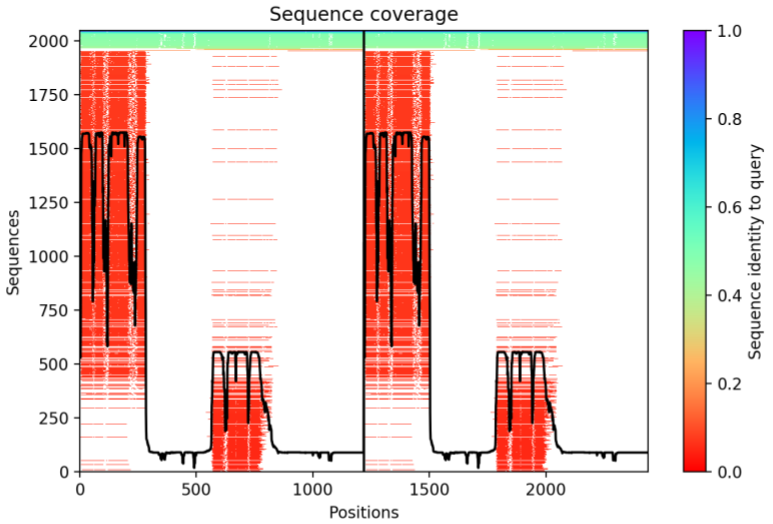

Sequence coverage
Sequences
Positions
Sequence identity to query
0
250
500
750
1000
1250
1500
1750
2000
0
500
1000
1500
2000
0.0
0.2
0.4
0.6
0.8
1.0


**C**

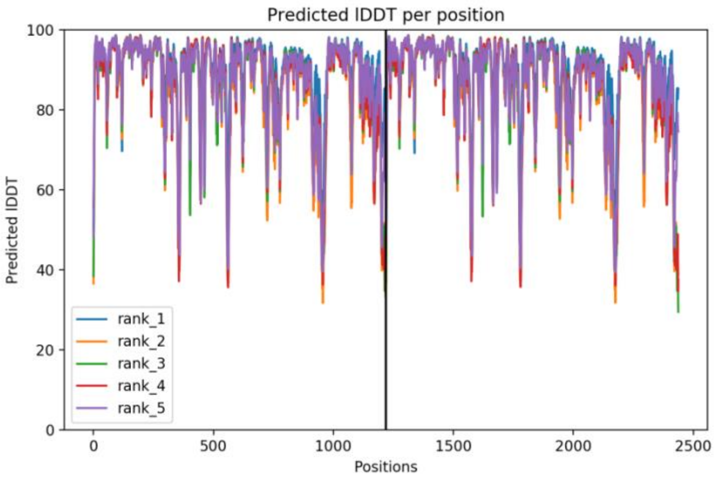


**D**

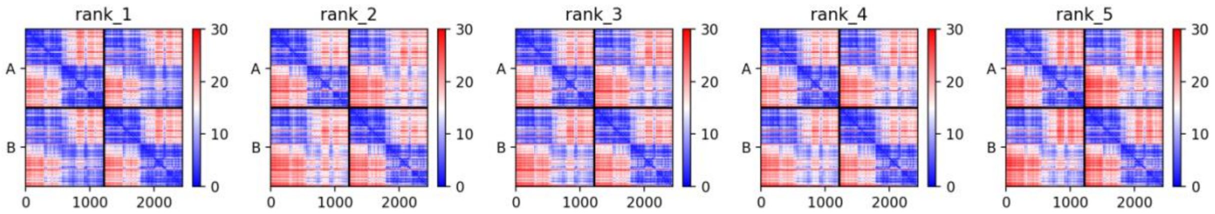


**Figure S15. AlphaFold prediction of *Ca*MCR. A** Superposition of five ColabFold *Ca*MCR dimers. B Sequence coverage of the MSA for the prediction. C plDDT score per position. D. predicted alignment error matrices (PAE) for the predictions. Each individual domain is predicted well (blue squares on the diagonal), the interaction of each domain with its dimer partner is also predicted well (off diagonal blue squares), but the orientations of unlike domains are less well predicted (red squares).
